# Citrullination of TDP-43 is a key post-translation modification associated with structural and functional changes and progressive pathology in TDP-43 mouse models and human proteinopathies

**DOI:** 10.1101/2025.02.28.639952

**Authors:** Christopher Saunders, Patricia Rocha-Rangel, Rohan Desai, Zainuddin Quadri, Haiying Lui, Jerry B. Hunt, Huimin Liang, Colin Rogers, Kevin Nash, Phoebe S. Tsoi, Erin L. Abner, Allan C. Ferreon, Josephine C. Ferreon, Christopher Norris, Vladimir N. Uversky, Peter T. Nelson, Jillian Cramer, Dale Chaput, Daniel C. Lee, Maj-Linda B. Selenica

**Author notes:** **Corresponding author:** Please direct all correspondence including post-publication to: Maj-Linda B Selenica, PhD Assistant Professor, Sanders-Brown Center on Aging Department of Molecular and Cellular Biochemistry College of Medicine, University of Kentucky 789 Todd Lee Jr. Building Lexington, Kentucky, 40536.

## Abstract

TAR DNA-binding protein 43 (TDP-43) pathology is associated with a spectrum of clinical dementias including limbic-predominant age-related TDP-43 encephalopathy neuropathological changes (LATE-NC). Post-translational modifications (PTM) are linked to TDP-43 toxic gain-of-function and cytoplasmic aggregation^1–3^. Phosphorylation remains the most investigated PTM and a standard criterion for determining pathology progression and clinical subclassification in TDP-43 proteinopathies^4–7^. However, full spectrum of PTMs on TDP-43 structure and biology remain unknown. Utilizing mass-spectrometry analysis we identified *citrullination* as a novel and irreversible “bona-fide” PTM of TDP-43 protein. We recognized peptidyl arginine deiminase 2 and 4 (PAD2 and PAD4) to mediate the conversion of arginine (R) to citrulline (citR) *in vitro* and demonstrated increased PAD2 and PAD4 expression and TDP-43 citrullination in a human wildtype TDP-43 mouse model (Tg (Thy1-TARDBP4). Transmission electron microscopy imaging analysis revealed citrullination induced vast structural changes while ThT analysis demonstrated altered aggregation kinetics of citrullinated (citR) TDP-43 protein. We further provided mechanistic evidence on reduced electrostatic and pi-pi interactions of citR TDP-43 Low Complexity Domain (LCD) with RNA, favoring liquid-solid phase separation and condensate formation. Generation and validation of several citR TDP-43 specific antibodies against several TDP-43 epitopes revealed epitope and domain-specific effects of citrullination on TDP-43 solubility *in vivo*. Importantly, we found distinct reactivities of citR TDP-43 antibodies shedding light into the contribution of epitope-specific properties of human citR TDP-43 to novel pathological citR TDP-43 assemblies in human brain tissue from LATE-NC, with or without comorbid Alzheimer’s disease neuropathologic changes (ADNC). These findings provided a unique look into the temporal citR TDP-43 signatures, and the potential clinical relevance associated with progression of pure LATE-NC and comorbid ADNC + LATE-NC. Collectively, these data reveal the existence of irreversible TDP-43 citrullination at targeted sites via induced PAD2/PAD4 activities, presenting a critical step in TDP-43 proteinopathy.

## Introduction

TDP-43 (transactive response DNA binding protein of 43 kDa) is a primarily nuclear-localized RNA/DNA binding protein associated with a spectrum of neurodegenerative diseases^1^. In recent years TDP-43 Neuropathologic Changes (NC), which encompass the clinical and pathological course of TDP-43 proteinopathy, were implicated in ∼50% of demented individuals 85 years or older in Limbic-Predominant Age-Related TDP-43 Encephalopathy (LATE)^4–7^. LATE-NC commonly co-exists with Alzheimer’s disease (ADNC), hippocampal sclerosis (HS), and Lewy body (LB) pathologies in advanced disease and age^4,6,8–11^. Abundant evidence supports the hypothesis that TDP-43 proteinopathy contributes to cognitive decline and dementia severity in these disorders^12,13^.

The amino acid sequence of TDP-43 comprises of functional domains that include N-terminal nuclear localization sequence (NLS, aa 82-98), two RNA recognition motifs (RRM1, aa 106-176 and RRM2, aa 191-262), and a low-complexity, intrinsically disordered C-terminal domain (LCD, aa 266-414). Post-translational modifications (PTM) enhance the protein flexibility^14^ and often provide unique functional characteristics, but they can also play a critical role in disease pathogenesis. In TDP-43 proteinopathy, the most commonly studied reversible PTMs — hyper-ubiquitination, acetylation, and phosphorylation — are often linked with altered homeostatic TDP-43 structure and biological function, rendering TDP-43 toxic gain-of-function in disease states^1–3,15^. To date, phosphorylated TDP-43 (pTDP-43) remains a key pathologic marker for disease diagnosis and categorization in several disorders including amyotrophic lateral sclerosis (ALS), frontotemporal lobar degeneration with TDP-43 (FTLD-TDP) and LATE-NC ^16–20^.

Citrullination (citR) is an irreversible PTM catalyzed by peptidyl arginine deiminases (PADs)^21^. Human *PADi* (gene) is found on chromosome 1p36.1, and encodes five PAD isozymes, PAD1 - 4 and 6, with a sequence identity of 50% and varying tissue distribution, anatomic localization and thereby disease association^22–25^ By replacing the imine of the positively charged guanidinium group on arginine with neutrally charged urea^26^, there is a net reduction from the positive-charge and isoelectric point of the arginine (+1, pI = 11.41) to a neutral charge (pI = 5.91). This irreversible conversion increases protein hydrophobicity and alters protein conformation and intramolecular/ intermolecular interactions^25,27,28^.

Thought to natively exist in an inactive form, PADs are Ca^2+^-dependent enzymes, binding to five vs. six Ca^2+^ions per molecule in their active site, respectively, indicating the role of Ca^2+^ as a molecular switch in PADs activation during inflammation and cellular stress^26,29,28,30,31^. Notably, PAD2 and PAD4 are primarily expressed in astrocytic and neuronal populations of CNS^32–35^. Both enzyme’s catalytic activity on citrullination of proteins, including histones and ribonuclear proteins (RNP), has been reported in various cancers and autoimmune diseases, while high levels of citrullinated peptides and histone H3/ H4 serve as autoantigens in rheumatoid arthritis^36,37^.

Several early studies have reported increased PAD2 and 4 expression in tissue from ADNC^38^, multiple sclerosis (MS)^39^, Parkinson’s disease (PD)^40^, while increased citrullinated GFAP was reported in AD brain tissue^41^. In a recent report increased PAD2 levels correlated with heightened levels in ALS^32^, collectively suggesting that aberrant citrullination of proteins is implicated in the onset and progression of neurologic diseases^32,33,42–46^. In contrast, accumulation of FET proteins associated with ALS (FUS, EWS, and TAF15) and hnRNPA1 were reduced in a HeLa cellular model following PAD4-mediated citrullination, suggesting that citrullination could promote protein solubility^47^. Hence, despite great biological and clinical potential, the role of citrullination on various PAD substrates and their potential molecular mechanisms across the dementia spectrum remains limited.

In the present study, we performed multisystem analyses including recombinant protein, mice models and human brain to determine whether TDP-43 undergoes irreversible citrullination leading to changes in TDP-43 protein structure and folding kinetics associated with TDP-43 pathology progression in neurodegenerative disease. We identified PAD2 and PAD4-mediated conversion of 11 arginine to citrulline across all domains of TDP-43 protein and developed novel, site-specific antibodies against citrullinated TDP-43 at several key epitopes. Notably, we identified the presence of citR TDP-43 in LATE-NC and AD + LATE-NC brain tissue, revealing disease and regional differences. Overall, we demonstrate PAD-mediated TDP-43 citrullination as a hallmark of TDP-43 pathology and propose a mechanism by which citrullination-driven structural changes impair TDP-43 function.

## Methods

### TDP-43 protein enzymatic *in vitro* citrullination

*In vitro* citrullination was performed by incubating 10 µM recombinant human full-length TDP-43, 1-414 aa (LSBio, #LS-G2316, WA, USA) with recombinant human 5.8 µM PAD4 or PAD2 proteins (Cayman, #10500, #10785, MI, USA). Reaction samples were incubated in citrullination buffer (50mM boric acid, 2mM calcium dichloride anhydrous, pH 8.0) at 37⁰C for 90 minutes, with agitation every 15 minutes. Samples were then quenched on ice for five minutes to stop all enzymatic activity. Modified and unmodified protein was used immediately for most recombinant cell-free studies described herein or stored as aliquots in -80⁰C for further biochemical analysis.

### Mass Spectrometry-based Proteomics

For the mass spectrometry analysis, full length TDP-43 protein undergoing PAD2 and PAD4 mediated citrullination were denatured in 4% β-mercaptoethanol, 95⁰C for 10 minutes, and separated by SDS-PAGE (Bio-Rad, #135BR, CA, USA). The gel was then stained with Coomassie (Bio-Rad, #1610786, CA, USA). Bands were excised and processed for LC-MS/MS analysis. Each gel piece was minced and de-stained before being reduced with dithiothreitol (DTT), alkylated with iodoacetamide (IAA), and digested with Trypsin/Lys-C overnight at 37°C. After digestion, peptides were extracted using 50/50 acetonitrile (ACN)/H_2_O/0.1% formic acid and dried in a vacuum concentrator. Dried peptide samples were resuspended in 98%H_2_O/2%ACN/0.1% formic acid for LC-MS/MS analysis.

Peptides were separated on a 50cm C18 reverse-phase UHPLC column (Thermo Fisher Scientific, MA, USA) using an Ultimate3000 UHPLC (Thermo Fisher Scientific, MA, USA) with a 60-minute gradient (4-40%ACN/0.1% formic acid) and analyzed on a hybrid quadrupole-Orbitrap mass spectrometer (Q Exactive Plus, Thermo Fisher Scientific, MA, USA). The mass spectrometer was operated in data-dependent acquisition mode, using a resolution of 70,000 m/z. The top 10 most abundant ions were selected for MS/MS, with an isolation window of 1.5 Da.

The raw data files were processed in MaxQuant (CoxLab, v2.6.7.0) and searched against the Uniprot *Homo sapiens* TDP-43 sequence database (UniProt ID: Q13148). Proteins were identified using a filtering criterion of 1% protein and peptide false discovery rate, with a peptide tolerance of 4.5 ppm. Search parameters included modification of cysteine by carbamidomethylation, and the variable modifications, methionine oxidation, arginine citrullination, as well as asparagine and glutamine deamidation. MS/MS spectra were also manually annotated to verify the sequence coverage of each citrulline containing peptide. Additionally, the conversion of arginine to citrulline increases the hydrophobicity of the peptide, resulting in a later retention time than its unmodified counterpart. This change in retention time also allows the differentiation between peptides containing citrullination, and those containing asparagine or glutamine deamidation^48^. Citrullination was considered only if they were reliably distinguished from deamidation, and they presented sufficient b- and y-fragment ion coverage surrounding the modified site. The most representative MS/MS spectra for each citrullinated peptide was selected for manual annotation and extracted ion chromatograms (XICs) were generated for each peptide identified to determine their retention times.

### Molecular weight shift by SDS-PAGE

Upon citrullination with PAD2 or PAD4 enzyme, modified and unmodified TDP-43 samples were denatured and separated by SDS-PAGE as described above, transferred to a PVDF membrane, and incubated with a total TDP-43 antibody (Proteintech Group Inc, #10782-2-AP, IL, USA). Modified (citrullinated) and unmodified TDP-43 were imaged via chemiluminescence. Molecular weight shift of citrullinated TDP-43 band was determined in ImageJ (NIH, version JAVA 1.8.0_322, ME, USA). Briefly, a standard log curve was fitted to the linear distance of each standard band measured from the largest molecular weight band (Fisher BioReagents^TM^, #BP3603-1, MA, USA). Values were plotted to interpolate the unknown molecular weight (mw). The difference between conditions determined Δmw value.

### In vitro TDP-43 turbidity assays

Following *in vitro* citrullination, 50µL of unmodified and citrullinated recombinant 10µM TDP-43 (1-414 aa) was added to a 96-well black sided, clear bottom plate (Corning Costar, #3603, NY, USA), in triplicates. Protein was incubated with 50uL of assembly buffer (20mM HEPES, 150mM sodium chloride, and 1mM DTT, pH 7.4) and in the presence of 250ng/µL yeast total RNA (Millipore Sigma, #10109223001, MO, USA), covered in optical PCR film. Turbidity assay was performed in a final concentration of 5 µM TDP-43 for 32 hours and citR TDP-43 for 160 hours. Absorbance at 395nm was registered at 2-minute intervals and 4x bright field images were taken every 2 hours with continuous orbital agitation at 37⁰C (800 cfm, Biotek Cytation5, #CYT5MFW). Spectra data were analyzed and graphed using GraphPad Prism.

### Phase separation and Thioflavin T fluorescence assay

Recombinant Low Complexity Domain of TDP-43 C-Terminal (LCD, 263-414 aa) and assembly imaging and analysis was performed as described with few modifications^49^. Briefly, 10, 20, and 40 µM LCD protein was citrullinated in 1 : 2 PAD4 molar ratio as described above. citR LCD was re-purified and PAD4 was separated via revers-phase HPLC following LCD purification methods^49^. Citrullinated and unmodified LCD was incubated in αβγ buffer (10mM sodium acetate, 10mM sodium phosphate, 10mM glycine, 200mM NaCl, pH 7.5) with 10 µM thioflavin T (ThT; Millipore Sigma, #T3516, MO, USA). 0. 31.25 or 250 ng/µL polyU RNA was added to induce phase separation. 10 µL sample volume were pipetted onto 35-mm glass bottom dishes (ibidi, Martinsried, Germany) and droplet formation was imaged using EVOS FL imaging system (Thermo Fisher, MA, USA). Fluorescent (GFP channel) and bright field images were taken at different timepoints to monitor LLPS droplet aging.

Quantitative analyses of fluorescent intensity in images were performed using NIS Elements-AR image analysis software (Nikon, RRID:SCR_014329, Tokyo, JPN). Images were analyzed by intensity threshold masking and pixel correlation to the binary area of individual objects within the field of view (FOV). Masking and thresholding values were kept consistent across all conditions. Values for individual object size, count, and field of view (FOV) binary area coverage were measured.

### Amyloid folding kinetic

Protein mixed in 100 µL citrullination buffer (TDP-43) or αβγ buffer (LCD) was first filtered through a 0.2 µm membrane and added in triplicates to a 96-well black sided, clear bottom plate (Corning Costar, #3603, NY, USA). The plate covered with optical PCR film were incubated in a multi-plate reader (Biotek Cytation5, #CYT5MFW, VT, USA) at set temperature of 37⁰C. ThT fluorescent spectra (excitation λ: 440/20, emission λ: 480/20) were read at 10-minute intervals with continuous orbital agitation (800 cfm). Spectra were analyzed using a logarithmic growth curve-fit analysis (GraphPad Prism and Microsoft Excel). The *t_1/2_* of each replicate curve was determined at 50% relative fluorescent unit (RFU) growth. The *t_lag_* of each replicate curve was determined by the linear intersection of *t_1/2_* and lag phase slope. *Ƭ*, or the inverse apparent rate constant 1/*k_app_*, was calculated at

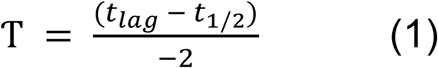

which were then used to determine best fit curves using the sigmoidal formula

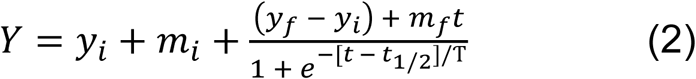

Residual plots were analyzed to determine fitness of curve^50^. All conditions were performed in triplicates and repeated in three independent experiments.

### Electron microscopy

For transmission electron microscopy (TEM) and scanning transmission electron microscopy (STEM), 300 mesh carbon coated copper grids (Electron Microscopy Sciences, CF300-CU, PA, USA) were washed with 10 µL Millipore water for 30 seconds. This was followed by a one-minute adsorption of 5 µL TDP-43 and citR TDP-43 samples alone or those incubated with yeast RNA as described above. Subsequent negative staining was performed with 10µL of 1% (w/v) filtered uranyl acetate (Electron Microscopy Sciences, #22400, PA, USA) for 30 seconds. Grids were capillary dried with laterally placed Kimwipes, then allowed to dry at least 4 hours before imaging. Sample grids were imaged for TEM (17,000 - 200,000x magnification), STEM, and TEM tomography using a FEI Talos F200X G2 (S)TEM (Thermo Fisher, MA, USA).

### TEM tomography and volume analysis

Tomography imaging was performed on the FEI Talos F200X G2 TEM. The sample grids were rotated around a horizontal axis with a range of -60⁰ to 60⁰ from horizontal zero, with images taken every 2⁰ for a total of 60 cross-sectional “slices”. Images were subsequently aligned and stacked using Inspect3D (Thermo Fisher, MA, USA) to produce binary data stacks.

Binary images were processed in ImageJ for frame-by-frame video production, (Avizo, Thermo Fisher) for mesh body rendering, and ChimeraX (v1.4, UCSF)^51,52^ and Fusion360 (Autodesk, version 16.6.1.2145, CA, USA) for body rendering. Volumetric dimensions, including surface area and volume, were generated by Avizo (Thermo Fisher). Branch size analysis for unmodified TDP-43 or citR TDP-43 was performed from each tomography binary stack by cropping individual branch ROIs, rotating to a vertical orientation, and orthogonally re-slicing to obtain a longitudinal cross-section stack along the z-axis. The intensity threshold mask equaled the binary value of the target branch. Ferret minimum diameter, branch area and volume were then analyzed with ImageJ software. ChimeraX is developed by the Resource for Biocomputing, Visualization, and Informatics at the University of California, San Francisco, with support from National Institutes of Health R01-GM129325 and the Office of Cyber Infrastructure and Computational Biology, National Institute of Allergy, and Infectious Diseases.

For video preparation, molecular surface volume renderings and analyses were created by processing the binary stack of image slices exported during tomography, conserving volumetric metadata. In brief, a binary image stack representing each condition was uploaded to ChimeraX (UCSF) and resolved with step 1 surface volume rendering. Renderings and its metadata were subsequently exported as an .STL file and modeled for 3D video in Fusion360 (Autodesk).

### Structural Alignment

Protein data bank nuclear magnetic resonance-derived structures of full length human TDP-43 (PDB ID: AF-Q13148-F1) were aligned in Swiss-PDB Viewer 4.0.1 (Swiss Institute of Bioinformatics, Lausanne, CH) and alpha fold using the magic fit function, resulting in an α-carbon RMSD of 15 Å or 5 Å (calculated for 64 atoms). Citrullinated arginine amino acids of each domain and relevant hydrogen bonds were revealed leaving the rest masked from view, then images were exported and rendered using Illustrator.

### Computational “pseudo-citrullination” disorder analysis

To model the potential effect of citrullination on TDP-43 residual and regional disorder propensity, all arginine residues (UniProt ID: Q13148) were computationally converted to glutamine (Q) (R→Q). The resulting “pseudo-citrullinated” sequence was utilized in subsequent bioinformatics studies. Per-residue disorder propensity of human TDP-43 protein (UniProt ID: Q13148) in non-modified and pseudo-citrullinated forms was evaluated by the Rapid Intrinsic Disorder Analysis Online (RIDAO) web platform^53^, which combines the outputs of residue disorder predictors, PONDR^®^ VSL2, and PONDR^®^ VLXT to generate the disorder profiles of query proteins. The disorder score was assigned to each residue, with a disorder score equal to or above 0.5 being considered as disordered and below 0.5 being predicted as ordered. Residues/regions with disorder scores between 0.15 and 0.5 were considered as ordered but flexible.

FuzDrop^54^ was used to predict spontaneous liquid-liquid phase separation and generated scoring system based on the protein sequence to identify the regions that promote this process. Protein with p_LLPS_ (probability of liquid-liquid phase separation) score of 0.60 or higher are identified as promoters of liquid-liquid phase separation^55^.

### Citrullinated TDP-43 antibody generation

The rabbit polyclonal antibodies were raised against TDP-43 citrullinated epitopes at position R83, R191, R165, R268, R275, R268/272, R293 (21^st^ Century Biochemicals, MA, US) utilizing a customized polyclonal antibody cross-affinity purification service. Briefly, rabbits were immunized with a citrullinated peptide for each corresponding citrulline epitope of human TDP-43 applying manufacture’s PVTV^TM^ platform. TDP-43 citrullinated antibodies were purified using affinity chromatography with modified and unmodified peptides following Q/C immunodepleting approaches. The cross-affinity purified citrullinated TDP-43 antibodies were then validated in recombinant enzymatic by western blot and immunohistochemistry utilizing brain tissue form transgenic TDP-43 mouse model and non-transgenic littermates as indicated below.

### Animals

Animal procedures were approved by the Institutional Animal Care and Use Committee of the University of Kentucky and performed in accordance with the eighth edition of the “Guide for the Care and Use of Laboratory Animals,” published by the National Academy of Science, the National Academies Press, Washington, DC^56^.

The human wildtype TDP-43 mouse model (wtTDP-43 Tg(Thy1-TARDBP4(TAR4 hemizygous line) was purchased by the Jackson Laboratory (The Jackson Laboratory, Stock #: 012836, Bar Harbor, ME, USA). TAR4/4 homozygous line was bred and maintained in the laboratory under approved IACUC protocols by University of Kentucky. Homozygous TAR4/4 mice (TAR4/4) express 2-fold expression of human wtTDP-43 and develop severe cytoplasmic TDP-43 pathology, abnormal limb reflex at 14 days and reduced motor performance by 18 days ^57,58^. Hemizygous TAR4 (TAR4) mice express 1.4-fold human wtTDP-43 transgene to the mouse gene and displays mild TDP-43 neuropathological profile with age^57,58^. In this study, we used 19-20 days old TAR4, TAR4/4 and age-matched littermates (n = 5 mice/genotype).

### Biochemical Fractionation and Western Blotting

Posterior cortex from TAR4 and TAR4/4 mice or non-transgenic littermates (Non-Tg), were fractionated as reported^59,60^. Briefly, human tissue was resuspended in 10% w/v modified RIPA buffer (50 mM Tris, 150 mM NaCl, 0.5% Triton x100, 1mM EDTA, 1mM EGTA, 1% sodium deoxycholate, 1% SDS), and mouse tissue was resuspended in 10% w/v RIPA buffer (50 mM Tris, 150 mM NaCl, 1% NP40, 0.05% sodium deoxycholate, 0.1% SDS), both with protease inhibitor cocktail (Sigma-Aldrich, MO, USA,#P8340) and phosphatase inhibitor cocktails II and III (Sigma-Aldrich, MO, USA, #P5726 and Cat #P0044). Tissue was homogenized manually followed by a brief (30 seconds) sonication pulse. Samples were ultracentrifuged at 38,000 x *g* for 30 minutes at 4⁰C and supernatant was referred to as RIPA-soluble brain homogenate. Protein concentration was determined by Pierce BCA protein assay (Thermo Fisher Scientific, #23225, MA, USA). Pellets were resuspended in urea buffer (7M urea, 2M thiourea, 4% CHAPS, and 30mM Tris; pH 8.5), sonicated and then ultracentrifuged at 100,000 x *g* for 30 minutes at 4⁰C and resulting supernatant was collected as Urea-soluble fraction. Protein concentration was determined by No-Stain Total Protein Assay via UV detection (Thermo Fisher Scientific, #A44717, MA, USA).

For SDS-PAGE, 10 μg of protein was denatured in 5x sample buffer (100mMTris pH 6.8, 4% SDS, 0.1% Bromophenol blue, 20% glycerol, 200mM β-mercaptoethanol) and run on 4-20% Tris-Glycine gel at 140V for 2 hours. Proteins were transferred to a PVDF membrane and then incubated on blotting-grade blocker in 0.7% TritonX-100 PBS (TBST, pH 7.4, + 0.7% Tween). The membrane was incubated overnight with the following citrullinated (citR) primary antibodies; citR83 TDP-43 for (NLS motif), citR165 (RRM1), citR191 (RRM2), citR275, citR268/272 (CTD) at 1:1000 dilution factor. Membrane was also probed for anti-human TDP-43 antibody (Sigma-Aldrich, #WH0023435M1, MO, USA), total (human + mouse) TDP-43 and phosphorylated S409/410 TDP-43 (Proteintech Group Inc, #10782-2-AP, #22309-1-AP, IL, USA, respectively). Following 3x TBST rinses, the membrane was incubated with HRP conjugated host-specific secondary antibody for 2 hours (Southern Biotech, anti-rb #4030-05, anti-ms #1071-05, AL, USA) and enhanced chemiluminescence (Thermo Scientific, #32106, MA, USA) was used for protein detection. Signal was normalized to GAPDH (Proteintech Group Inc, #HRP-60004, IL, USA) or No-Stain total protein for RIPA and Urea fractions, respectively.

### Immunoprecipitation - Mass spectrometry

Immunoprecipitation was facilitated using Protein A trisacryl resin beads (#P3391, Millipore-Sigma, MO, USA). Brains from TAR4/4 and non-Tg control mice were prepared using the previously described homogenization protocol. Resulting brain homogenate (input) was immunoprecipitated using 4 µg of TDP-43-antibody (C-terminal TDP-43; Proteintech Group Inc, #12892-1-AP, IL, USA) that was conjugated to resin beads. Aliquots of brain homogenate were standardized to 1.5 mg, then combined with the conjugated beads to incubate overnight at 4°C. After overnight incubation, the beads were washed twice with buffer (RIPA + inhibitors) and once with water. The freshly washed beads were then incubated and rinsed with elution buffer (0.1 M glycine, pH 2.0) twice. Each ‘elution’ was collected and combined into one tube, and neutralization buffer (1M Tris, pH 8.0) was added to the final sample. Once neutralized, the samples were passed through concentrator columns (Thermo Scientific, #88512, MA, USA) to produce the final immunoprecipitated product for mass spec. IP’d samples at pH 7.0-8.0 and input brain homogenate were denatured in 5x sample buffer with 4% β-mercaptoethanol at 95°C for 10 minutes. Membrane was probed against a total TDP-43 antibody (Proteintech Group Inc, #10782- 2-AP, IL, USA).

For LC-MS/MS analysis, the IP eluate was digested, and peptides were extracted using 50/50 acetonitrile (ACN)/H_2_O/0.1% formic acid) and dried in a vacuum concentrator. Dried peptide samples were resuspended in 98% H_2_O / 2% CAN / 0.1% formic acid for LC-MS/MS analysis following similar protocol as described for the recombinant protein.

### Immunohistochemistry animal tissue staining and analysis

For citR TDP-43 antibody characterization and the study, six sections per mouse (n = 5 / group), collected 150 μm apart, underwent immunohistochemical analysis as described previously^61^. Briefly, half hemisphere of the mouse brain was fixed in 4% PFA fixed and underwent sequential dehydration in 10%, 20%, and 30% sucrose buffers for 24 hours. Brain sections of 25 µm thickness were stored in DPBS + sodium azide at 4°C until immunohistochemistry assays were performed following previously published protocols^59,62^. First, all citR TDP-43 antibodies were validated in a log-fold antibody serial dilution; tissue was incubated in primary antibody dilutions ranging from 1:30,000 to 1:300, including incubation with secondary antibody only (no primary antibody control).

Optimal citR TDP-43 antibody concentrations were determined above, however, tissue stained with citR191 and citR268/272 TDP-43 antibodies were first treated with 70% formic acid for 15 minutes at room temperature and rinsed with 2x DI-water. Endogenous peroxidase was blocked (10% methanol, 10% H_2_O_2_ in PBS). Tissue was permeabilized (with 0.2% lysine, 1% Triton X-100 in PBS) and incubated overnight in primary antibodies; citrullinated R83 (NLS), R191 (RRM2), and R268/272 (CTD) TDP-43 at 1:3000, 1:1000 and 1:1000 dilution factor, respectively. PAD2 and PAD4 primary antibodies were purchased from Proteintech, US (#12110-1-AP and 17373-1-AP) and diluted at 1:10000 vs. 1:1000, respectively. Sections were incubated with anti-rabbit biotinylated secondary antibody and enzymatic conjugation with Vectastain® Elite® ABC kit was performed for 1 hour (Vector Laboratories, #PK-6100, Burlingame, CA, USA; cat # PK-6100). Finally, sections were developed using 0.05% diaminobenzidine, 0.5% Ni^2+^ and 0.03% H_2_O_2_, mounted onto slides, dehydrated, and coverslipped. Tissue incubated without citR TDP-43 primary antibodies (no primary control) were included to confirm antibody specificity *in vivo*.

Imaging was performed using a Zeiss Mirax150 digital scanning microscope (Carl Zeiss MicroImaging, GmbH Clinical, 07740 Jena, GER). For regional quantification, parameters such as hue, saturation, and intensity were used for each image field. Object segmentation thresholds were established from images of high and low levels of staining to identify positive staining over the intensity levels in the study. These limits were held constant across this study as reported by Gordon and colleagues^61^. The edge of the layer was defined as the furthest visible cells continuous with the layer (Carl Zeiss MicroImaging, GmbH Clinical, 07740 Jena, GER).

### Human subjects

Post-mortem brain tissues were provided by the University of Kentucky Alzheimer’s Disease Research Center (UK-ADRC) and the University of California Irvine Alzheimer’s Disease Research Center (UCI-ADRC) brain banks. All cases were assessed systematically for pathology, genotyping, and biochemical analyses from UCI=ADRC and UK-ADRC Neuropathology Core facilities ^63–65^. LATE-NC staging was based on the published consensus recommendations: LATE-NC *stage 1*: pTDP-43 pathology was limited to amygdala; *stage 2:* pTDP-43 pathology was extended into the cornu armmonis (CA1 and 3) neuronal layers and dentate gyrus of hippocampus (61–64). Pure LATE-NC was as defined previously^66^. ADNC diagnosis followed National Institute of Aging - Alzheimer’s Association consensus criteria and we included Braak NFT stages IV-V in the current study to indicate the presence of ADNC^67^. Autopsies were performed after consent in accordance with institutional review boards. Frozen human brain tissue or brain sections were processed for SDS-PAGE and Western blotting, or immunohistochemical (IHC)/ immunofluorescence (IF) labeling procedures, respectively, as described below.

### Human Brain Immunohistochemistry

*For formalin-fixed, paraffin-embedded (FFPE) sections*: Slide-mounted sections (8-10 µm thickness) from LATE-NC stage 1 (amygdala), stage 2 (hippocampus), and healthy control were deparaffinized in xylene, followed by a series of decreasing concentrations of ethanol. Both PFA-fixed and FFPE sections were briefly rinsed in water for 5 minutes before heat-induced antigen retrieval; slides were microwaved in Diva Decloaker buffer, pH 6.0 (Biocare Medical, CA, USA) for six minutes. Endogenous peroxidase was blocked (3% H_2_O_2_ in methanol for 30 minutes) followed by rinse in 1 X TBS-Triton (0.1% Triton X-100). Sections were blocked for 1 hour with 5% serum prepared in TBS-Triton and 3% BSA buffer (pS409/410 TDP-43, in rabbit serum, and all citR TDP-43 antibodies, in goat serum). Primary citR and pS409/410 TDP-43 antibodies were diluted in blocking buffer (1X TBS-Triton and BSA) and incubated at 4°C overnight. Slides were rinsed in TBS-Triton prior to incubation with the corresponding biotinylated secondary antibody. After 3 x TBS-Triton rinse, slides were incubated for 1 hour with the Vectastain® Elite® ABC kit (Vector Laboratories, #PK6100, CA, USA) and developed using ImmPACT NovaRED (Vector Laboratories, #SK-4805, CA, USA). Hematoxylin QS was used for nuclei counterstaining (Vector Laboratories, #H-3404-100, CA, USA) followed by cover slipping with Epredia mounting medium (Fisher Scientific, #22-110-610, PA, USA).

*For free-floating tissue sections*: the same protocols were used for human as the mouse tissue. Sections were mounted onto slides (two sections/brain) and dried overnight then subsequently placed into 37 °C oven for an additional 24 hours.

### Immunofluorescence and confocal imaging

TDP-43 immunohistochemistry was performed using pTDP-43 pS409/410 (1D3 clone, Biolegend, #829901, CA, USA) and citR268/272 TDP-43 to determine spatial and conformer specificity on selected human brain tissue. Secondary Alexa Fluor antibodies conjugated to anti-rabbit Alexa Fluor^TM^488 and anti-rat Alexa Fluor^TM^594 (Thermo Fischer Scientific, Carlsbad, CA, USA) were used. DAPI (4′,6 -diamidino-2-phenylindole) was used to visualize cell nuclei. Images were obtained with a Ti2 Nikon A1 confocal microscope (Nikon, Tokyo, Japan) using Nikon Plan Apo Lambda 20X and 40X lens objectives with NyQuist optical zoom enhancement. Z-stacks for each fluorophore were obtained by large-image stitch resonant scans using four sequential channels and corresponded to an average of eight frames at 1024 px^2^. Images were processed and analyzed using NiS Elements-AR software, with flattened and volume Max Intensity Projections created from all z-planes. Co-localization between fluorophores were analyzed by size, threshold, circularity, and ROI inclusion parameters; all acquisition — including laser, z-step, and camera setup — and analysis settings were kept constant across images.

### Citrullination profiling in human tissue and analysis

Human brain section imaging was performed using a Zeiss Mirax150 digital scanning microscope and as described above (Carl Zeiss MicroImaging, GmbH Clinical, 07740 Jena, GER). Regional quantification, parameters and object thresholding was also performed similar to the animal protocols. Labeling intensities for pTDP-43 and citR TDP-43 were quantified within regions of interest (ROI, n =7) for each brain tissue, including amygdala, hippocampus, and frontal cortex. Healthy controls and AD patients exhibiting TDP-43 pathology without LATE diagnosis ^(61–64)^ was excluded from analysis. Technical artifacts (tissue folds, etc.) present were also excluded to avoid false positive signal. Positive immunoreactivity for citR TDP-43 was expressed as percent area intensity per ROI.

### Statistical analysis

Statistical analyses were performed using paired Student’s *t*-test, one-way or two-way ANOVA as indicated in each figure, followed by *post hoc* multiple comparisons test, using GraphPad Prism (GraphPad Software, LLC, v10.1.2, Boston, MA, USA, www.graphpad.com). Values are represented as the standard error of the mean (SEM) unless stated otherwise. P values were considered statistically significant as followed: *p < 0.05, **p < 0.01, ***p < 0.001, ****p < 0.0001. The average percent positive area values measured by immunohistochemistry were normalized to the non-transgenic or healthy controls. Protein levels measured by western blot were normalized to GAPDH or no-stain protein, then graphed as fold change.

## Results

### Mass-spectrometry identification of TDP-43 citrullination

Modification of an arginine residue to citrulline occurs in low abundance in physiological systems and contributes only to a small 0.98 Da mass change^68^, previously proven to challenge the identification of citrullination by mass spectrometry-based proteomics. However, recent improvements in UHPLC have allowed for further chromatographic separation of citrullinated peptides and their unmodified counterparts^69^.

In this study, we sought to identify whether TDP-43 protein undergoes PAD - mediated arginine to citrulline modifications (Fig. 1a). To ensure accurate identification of modified arginine, we first performed PAD2 and PAD4 directed enzymatic reaction of recombinant TDP-43 and histone 3 (H3) protein, the latter representing a known PAD substrate^70,71^. This resulted in citrullination of H3 (citR H3) identified by western blot assay and using a specific citR H3 antibody (Fig. 1b). SDS-PAGE demonstrated a shift in TDP-43 molecular weight (MW) following enzymatic reaction with PAD2 and PAD4 protein (Fig. 1c, d), and calculations of band - distance from unmodified TDP-43 revealed an MW increase by 3.17 vs. 3.42 kDa, respectively (Fig. 1e).

**Figure 1.**
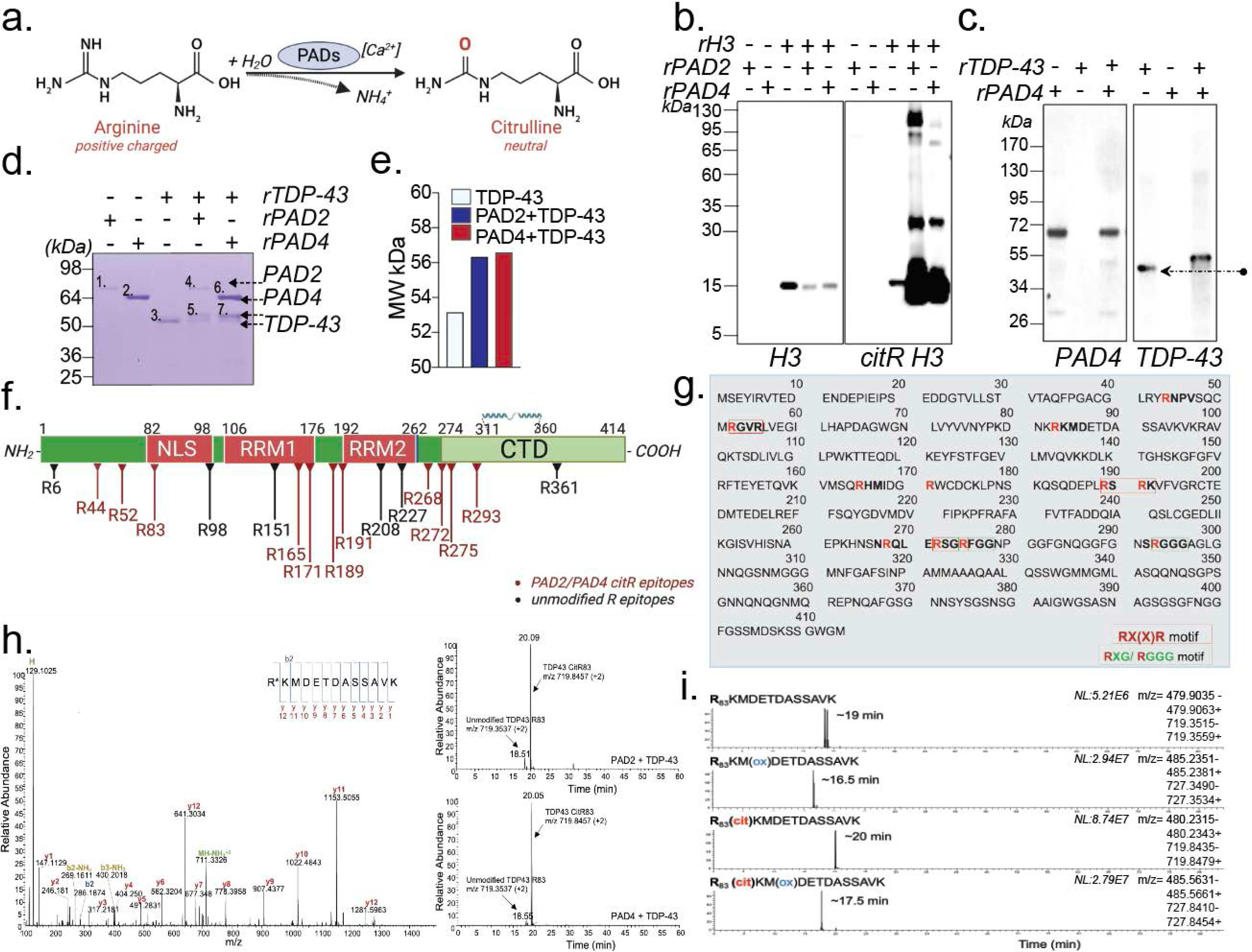
LC–MS/MS identification of citrullinated residues on recombinant human TDP-43 protein. a, The enzymatic reaction mediated by PADs converts arginine to citrulline. b, c, Western blot images of PAD2 and PAD4-mediated citrullination of H3 (b) and TDP-43 (c). In (c) arrows indicate the observed shift in TDP-43 molecular weight. d, Coomassie staining of PAD2 (bands 1, 4), PAD4 (bands 2, 6), unmodified TDP-43 (band 3), PAD2-mediated citR TDP-43 (band 5), and PAD4-mediated citR TDP-43 (band 7). e, Bar diagrams showing the MW shift (kDa) of TDP-43 protein following citrullination. f, Schematic representation of arginine epitopes positioned in the human TDP-43 protein sequence. Position of the 11 citrullinated arginine epitopes are indicated in red. g, Six out of eleven arginine epitopes susceptible to citrullination laid within common RX(X)R (red box) or RXG/RGGG (green box) motifs in TDP-43 sequence. h, MS/MS spectrum showing b- and y-ion coverage of modified citR83 peptide and extracted ion chromatograms (XICs) showed abundance and retention time of unmodified R83 *vs.* citR83 with PAD2 and PAD4 treatment (intact peptide monoisotopic m/z (+2) 719.8457, (+3) 480.2329). i, Retention time peaks for TDP-43 peptides surrounding the unmodified or modified R83 epitope and base peak m/z undergoing methionine (Met85) oxidation, citrullination or both showed reliable time separation.

To identify possible modifications, bands from unmodified and modified TDP-43 underwent in-depth analysis using liquid chromatography tandem mass-spectrometry (LC-MS/MS). Given that citrullination eliminates the positive charge of arginine under acidic LC-MS/MS conditions and often results in delayed peptide elution times^72^, we applied several factors and stringent separation techniques (described in Material and Methods). This was followed by rigorous data filtering using MAXIQuant and manual verification of each spectra that resulted in high degree of confidence to identification of citrullination at eleven out of twenty arginine residues across the TDP-43 protein sequence (Fig. 1f, Supplementary Table, localization score and PEP value). We further recognized the R*X*(*X*)R (Fig. 1g, red box), R*X*G or RGGG sequences (Fig. 1g, green box) as preferred TDP-43 recognition motifs for PAD2 and PAD4 activity. The extracted ion chromatogram (XIC) for each identified citrullinated peptide and the presence of b- and y-fragment ion coverage surrounding the modified site is shown for PAD4 and PAD2 modified (citR) *vs*. unmodified R83 (Fig. 1h) and R165, R191, R268/272 and R293 residues (Supplement Fig. 1a - l). We showed here the spectra and elution peaks for citrullinated R83 peptides, eluted at ∼ 20.5 min was separated from the unmodified arginine containing peptides, eluted at ∼18.5 min. The manual verification also allowed differentiation of asparagine and glutamine deamidation *vs.* arginine citrullination (*aka* deamination), as deamidation cannot be differentiated only by a shift in mass^72,73^. Lastly, to confidently confirm detection of citR TDP-43 *vs.* unmodified protein, we compared XIC spectra for oxidation of methionine at nearby residue (Met85_ox_), or in combination with citrullination at R83 position. We demonstrated elution times for Met85_ox_ at 16.5 min and distinct from citR83 + Met_ox_ peptides eluted at 17.5 min (Fig. 1i). Again, we robustly identified the citrullinated R83 epitope eluted at ∼20 min vs. unmodified peptides at ∼19 min as shown in Fig. 1i.

Together, our findings provide the first high-confidence evidence on the full profile of TDP-43 citrullination profile identified by mass spectrometry analysis. Given the position of citR epitopes within each functional domain of TDP-43 protein, we aimed herein to investigate their biological impact across multiple system and models.

### Structural characteristics of citrullinated TDP-43

To determine whether citrullination alters TDP-43 amyloid-like aggregation, we incubated recombinant full length human unmodified and citrullinated TDP-43 (citR TDP-43) in citrullination buffer for 160 hours under aggregation conditions as previously reported^74–77^. Samples underwent Transmission Electron Microscopy (TEM), which revealed vast structural changes (Fig. 2). TEM images revealed amyloid-like, large amorphous aggregate structure of the unmodified TDP-43 (Fig. 2a, 17,000 – 200,000 magnification range). Interestingly, citR TDP-43 protein exhibited smaller and granular assemblies, structural features that vastly distinguish it from the amorphous aggregates of the unmodified TDP-43 (Fig. 2b, 17,000 – 200,000 magnification range). Next, we performed Scanning Transmission Electron Microscope (STEM), a technique that utilizes fine 0.05 – 0.2 nm spot size electron beam to generate a detailed structural surface mapping^78^. STEM scanning visually demonstrated the disperse granularity of the citR TDP-43 external structure and branch properties compared to the amorphous TDP-43 structure (Fig. 2c and d, insets, Fig. 2f and g). Consistent with TEM images, the STEM analysis revealed significant reduction of citR TDP-43 conformers volume (Fig. 2h, 237-fold, p < 0.001, unpaired t-test), and area (Fig. 2i, 19-fold, p < 0.001, unpaired t-test) compared to the TDP-43 protein aggregate. Further, the cross-sectional analysis of the individual branches obtained by TEM tomography revealed other morphological details of the citR TDP-43 conformers, consisting of significantly smaller branch diameter as compared to the unmodified TDP-43 branch (Fig. 2j, 26.38 ± 1.164 nm *vs.* 101.1 ± 3.441 nm, ****p < 0.0001) and area (Fig. 2k, 1471 ± 39.39 nm^2^ *vs.* 18541 ± 1283 nm^2^, ****p < 0.0001), suggesting a smaller and novel citR TDP-43 assembly.

**Figure 2.**
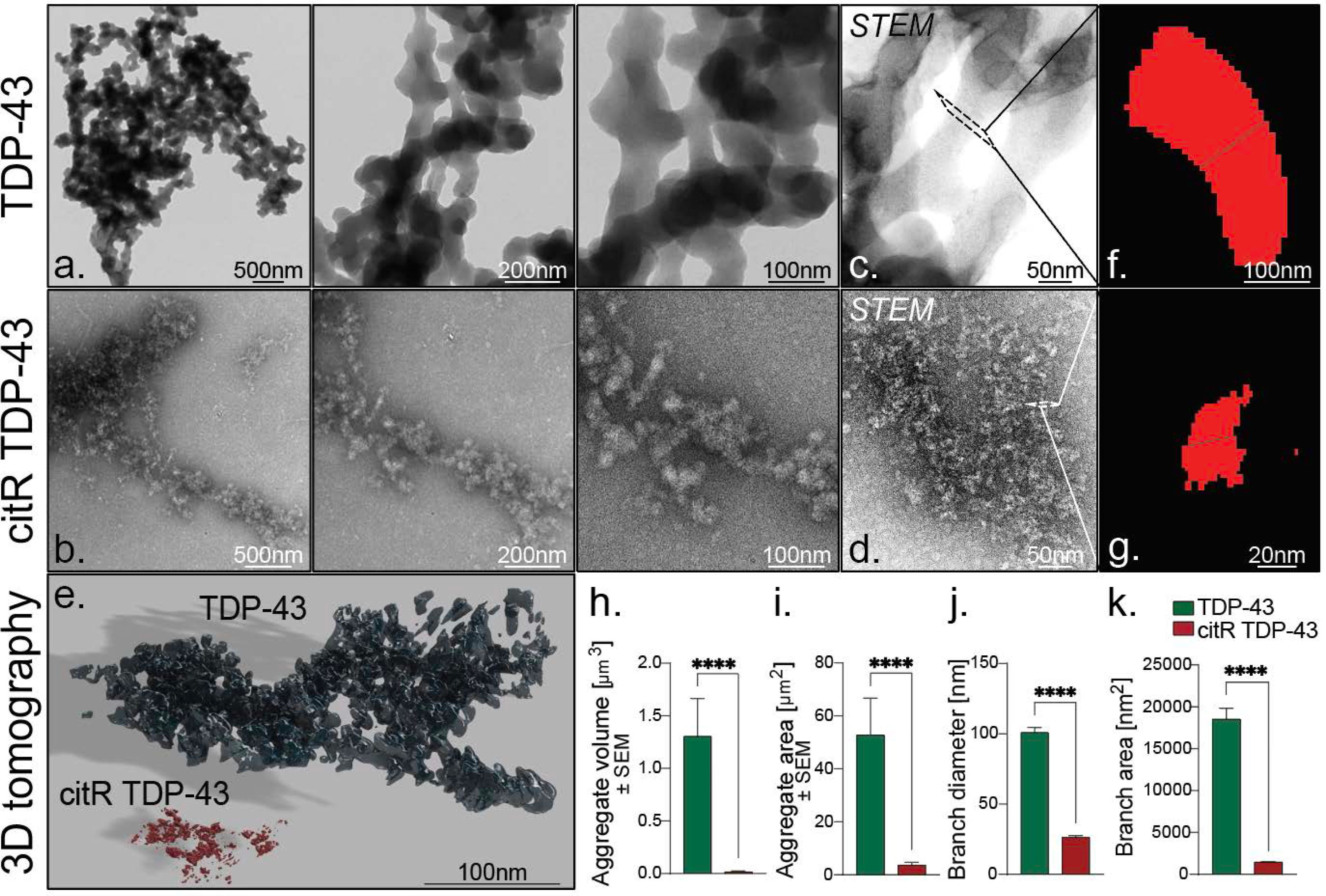
Citrullination of TDP-43 exposed the unfolding of amyloid amorphous aggregates and the morphological properties of citR TDP-43 protein conformers. Representative TEM and STEM images of a, c, unmodified TDP-43 amorphous aggregates and b, d, citR TDP-43 conformers at different magnifications following 5-day incubation. e, Representative 3D surface tomography renderings of both unmodified and citrullinated TDP-43 proteins. f, g, Representative cross-sectional branch image of unmodified TDP-43 vs. citR TDP-43 sectional surface, respectively. h, i, Analysis of aggregate volume (µm^3^) and area (µm^2^, and j, k, minimum branch diameter and branch area, determined by the mean of cross-sections across length of a branch. Values yielded from 3D tomography analysis, 3 ROI/ sample (n = 3). Student unpaired t-test, ****p < 0.0001. Scale bar is indicated in each image. Experiments were independently repeated at least three times.

Next, we performed TEM tomography and generated tilt-series of 2D projection images into a complete three-dimensional (3D) structural reconstruction^79^ for both unmodified TDP-43 and citR TDP-43 (Supplementary Video 1 and 2). The combined 3D tomography rendering of TDP-43 and citR TDP-43 proteins, confirmed the vast structural difference in the citR TDP-43 folded species (Fig. 2e and Supplementary Video 3).

The techniques and detailed analysis provided here uncover the structural properties that define the citR TDP-43 conformer and provide the basis for further investigation on the implications of such modification on TDP-43 interactions with target proteins and RNA.

### Citrullination impairs TDP-43 interactions with RNA revealing the biophysical characteristics of citR conformers

Several studies have suggested that TDP-43 physical interactions with (polyU) RNA are necessary not only for maintaining TDP-43 functional role in RNA splicing and metabolism^80,81^, but also halting its aggregation and regulating protein homeostasis and turnover^82–85^. To investigate the long-term effects of citrullination on TDP-43 interactions with RNA, we incubated unmodified or citR TDP-43 with yeast RNA (200ng/µl) for 160 hours. TEM imaging confirmed the amorphous structures of the unmodified TDP-43 aggregates (Fig. 3a), and the distinct granular structure of citR TDP-43 conformers (Fig. 3b). As expected, addition of yeast RNA interfered with TDP-43 protein aggregation and resolved the amorphous aggregates into smaller TDP-43 condensates (Fig. 3c, 3d 94,000x magnified). Surprisingly, we there were no morphological differences between citR TDP-43 protein alone (-RNA, Fig. 3b) and when incubated with RNA (Fig. 3e, 3f, 94,000x magnified images). Similar morphologies were also observed between unmodified TDP-43 incubated with RNA (+ RNA) and citR TDP-43 - RNA samples (Fig. 3c and 3b, respectively). Our data thus far suggested that citrullination of the arginine residues disrupted multivalent connections between TDP-43 protein and nucleic acid, necessary for regulating protein viscosity vs. elasticity properties^86^. The exact biophysical nature of citR TDP-43 : RNA interactions was not explored here but warrants future investigations.

**Figure 3.**
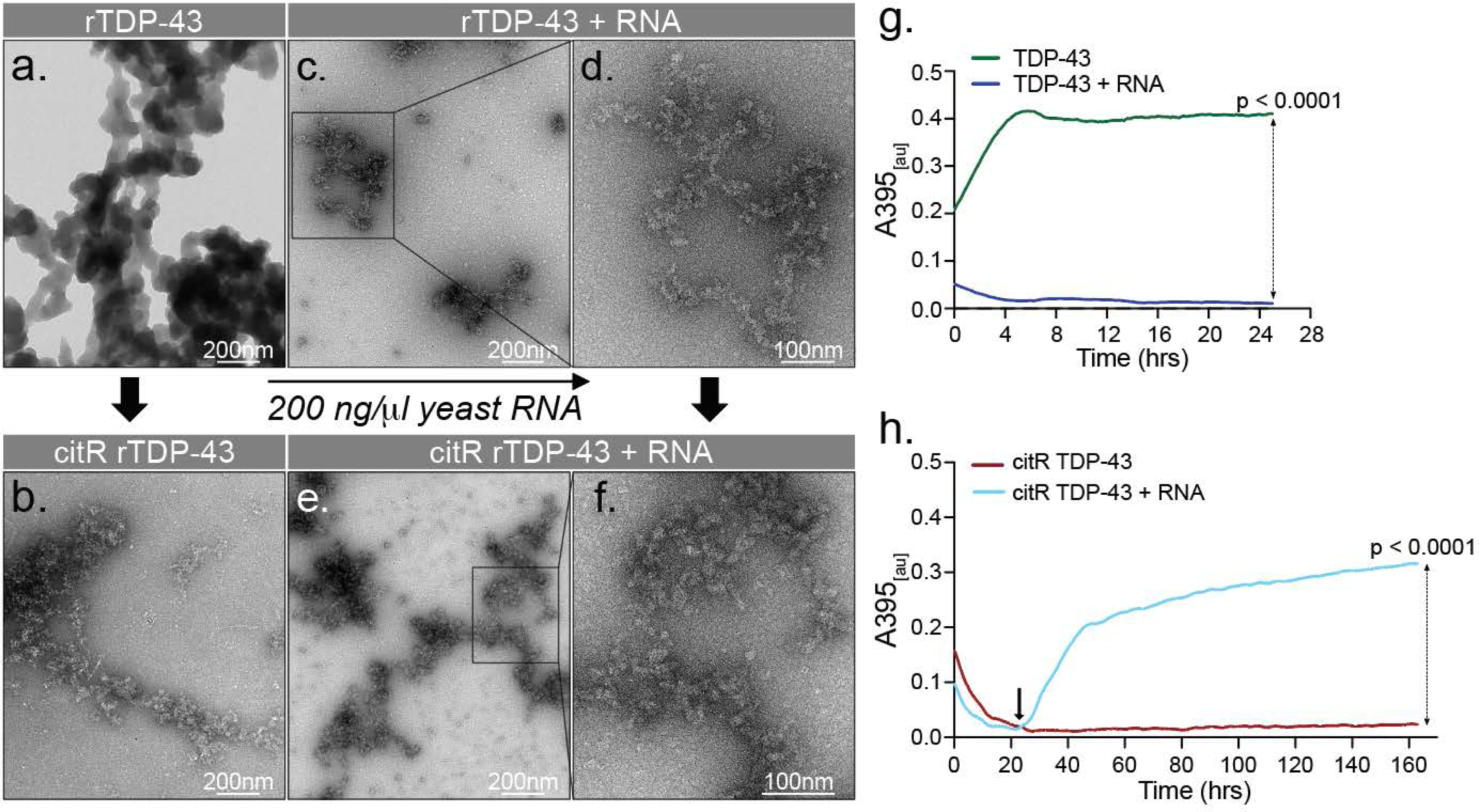
Citrullination of TDP-43 impairs protein – RNA interactions yielding citR TDP-43 condensates. a, b, TEM images of unmodified TDP-43 amorphous aggregates and citR TDP-43 structures following 5-day incubation. c, d, TEM images of unmodified TDP-43 aggregate and e, f, citR TDP-43 following 5-day incubation with 200ng/µl yeast RNA. Magnified TEM images of each structure (box, d, f). Scale bar is indicated in each image. g, Turbidity measured at absorbance wavelength of 395nm of the unmodified TDP-43 and h, citR TDP-43 in the absence and presence of yeast RNA. Arrow in (h) indicates the precipitation time point. Data are shown as the baseline-corrected means, n = 3 and repeated in at least three independent experiments. Student paired t-test, two-tailed, **** p < 0.0001.

To further analyze citR impact on TDP-43 protein biophysical properties and viscosity we performed turbidity assay^77^. We incubated unmodified and citR TDP-43 with yeast RNA (200 ng/µl) and absorbance was measured at (A)395 nm for 24 hours (Fig. 3g). Kinetic analysis showed that RNA significantly reduced unmodified TDP-43 absorbance indicating reduced protein viscosity (Fig. 3g, p < 0.0001). The citR TDP-43 protein alone also displayed low absorbance for up to 160 hr., suggesting a low viscosity state (Fig. 3h, dark red line). Intriguingly, citR TDP-43 + RNA sample maintained a low protein viscosity state in the first 24 hours (Fig. 3h, light blue line), followed by a sudden and significant increase in viscosity between 24 - 48 hr. time point, which slowly plateaued at 160-hour interval (Fig. 3h, arrow, p < 0.0001). Overall, this finding suggests that citrullination facilitates TDP-43 protein disaggregation but interrupts the physiological interactions with RNA, a process that results in induced protein viscosity *in vitro*.

### Computational simulations reveal citrullination-induced TDP-43 regional disorder propensities

To understand the impact of citrullination on TDP-43 aggregation and intrinsic disorder propensity, we conducted bioinformatics analyses of the full-length human TDP-43 protein (UniProt ID: Q13148). There are no natural codons of citrulline therefore, we computationally modeled “pseudo-citrullination” by converting all 20 arginine (R) residues present in TDP-43 to glutamine (Q) (R → Q) (Fig. 4a). We hypothesized that total R → Q “pseudo-citrullination” will affect the TDP-43 disorder state, as Q is not only a close citrulline-mimic but also one of the essential disorder-promoting residues from the following series: D < M < K< R < S < Q < P < E ^87,88^. We employed the RIDAO platform and included several individual disorder predictors to predict TDP-43 disorder profile driven by “pseudo-citrullination”. The disorder profile generated by PONDR^®^ VSL2 analysis demonstrated that “pseudo-citrullination” increased the local disordered propensity (disorder scores above 0.5) of the residues within the 150-220 aa region of TDP-43 protein (Fig. 4a, red dashed line), with a peak around the 180 aa (linker sequence between RRM1 and RRM2 regions). FuzDrop analysis^54,89^ predicted the liquid-liquid phase separation probability (LLPS) of sequences affected by “pseudo-citrullination” and revealed the formation of amylogenic “hot-spot” sequences within TDP-43 protein (Fig. 4b, c). Interestingly, “hot-spot” regions are predicted to drive the irreversible maturation of the liquid-liquid droplet to liquid-solid state^55,89^. As TDP-43 is characterized by high intrinsic LLPS potential, we determined TDP-43 residue-based droplet promoting probability (p_LLPS_) of 0.8981, verifying the sequence’ s tendency to undergo spontaneous liquid-liquid phase separation and droplet formation. FuzDrop analysis identified aggregation “hot-spots” at 251-255 (KGISV) and 317-355 (SINPAMMAAAQAALQSSWGMMGMLASQQNQSGPSGNNQN) residues within the TDP-43 droplet-promoting region (DPR, residues 251-414) (Fig. 4b). Notably, we found that TDP-43 R→Q “pseudo-citrullination” increased the overall p_LLPS_ to 0.9456 and identified three additional “hot-spots” regions within the DPR region spanning the 251-322 aa and 332-414 aa segments, including threw new “hot-spot” regions; the 321-330 aa residues of the amyloid core region (Fig. 4c), the linker sequence at residues 180-191 aa (SKQSQDEPLRSR), and the expended “hot spot” region between 251-264 aa residues in RRM2 (KGISVHISNAEPKH, boxed, Fig. 4c). To note, the predicted TDP-43 R → Q189 and R → Q191 residues within the “hot-spot” regions were “de facto” citrullinated by PAD2/ PAD4 in our LC-MS analysis (indicated in bold green). We identified three newly formed “hot-spots” within the CTD that included 310-322aa (GMNFGAFSINPAM), 332-342 aa (SSWGMMGMLAS), and 352-365 aa residues (NNQNQGNMQREPNQ (Fig. 4c).

**Figure 4.**
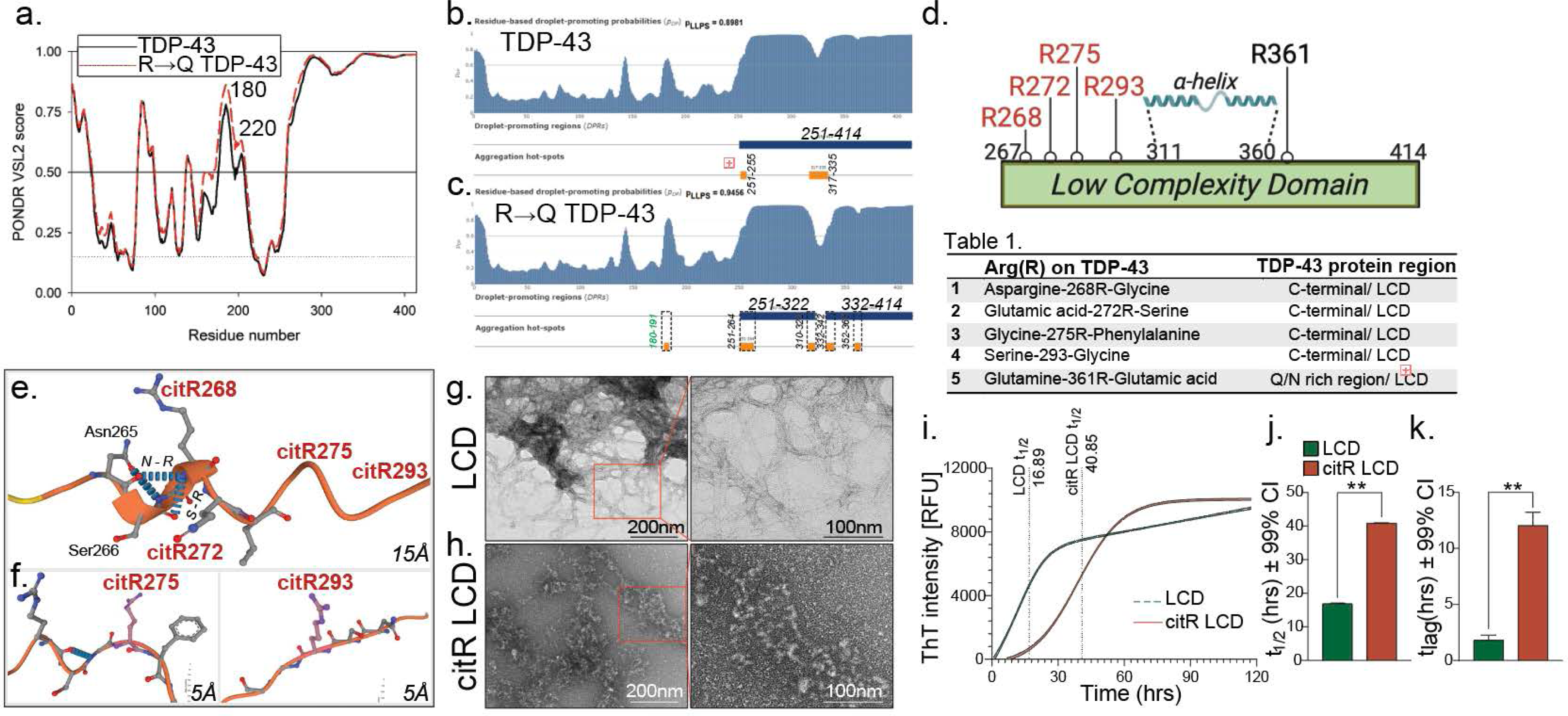
Characterization of citR TDP-43 intrinsic disorder and aggregation kinetic propensities. a, Bioinformatics analysis reveal the effect of TDP-43 pseudo-citrullination on protein disorder state. Changes in the intrinsic disorder state of human TDP-43 protein (black curve) following stimulated R→Q pseudo-citrullination (red curve) by PONDR^®^ VSL2 algorithm. b, FuzDrop algorithm predicted LLPS forming properties in TDP-43 (orange boxes) and c, simulated R→Q “pseudo-citrullination” introduced novel aggregation “hot-spot” regions/residues of TDP-43 (dashed orange boxes). Confirmed R residues citrullinated by PADs are indicated in green. Droplet-promoting regions (DPRs) are defined as 10 consecutive residues pDP ≥ 0.50. d, Schematic representation of citrullinated arginine epitopes within low complexity domain / LCD (aa 267-414 aa). e, f, Alpha-fold renderings of C-terminal TDP-43 (PDB ID:7KWZ) at R268, 272, 275 and 293 shown at 15Å and R275 and 293 at 5Å. g, h, Representative TEM images of aggregated unmodified and citR TDP-43 LCD at day 5 and magnified images (box). i, Relative fluorescence units (RFU) ThT intensity of 40 µM unmodified and citR LCD in the presence of 250 ng/µl RNA, revealing a delay in nucleation- and elongation-phase, calculated by *t_1/2_* and *t_lag_* values and j, k, Graphed *t_1/2_* and *t_lag_* values, respectively, at ± 99% CI (**p < 0.01), from non-linear curve-fit.

Overall, this analysis offered insights into the complex behavior of TDP-43 protein following modification of arginine to citrulline, impacting its structural flexibility (TEM) and physiological ⍺-helical LLPS dynamics of TDP-43 LCD domain (FuzDrop). We believe that these findings also facilitate interpretation of potential TDP-43 functional relationships following citrullination in physiology and disease.

### Citrullination causes significant structural and kinetic changes and delays TDP-43 LCD nucleation phase

Considering the impact of “pseudo-citrullination” on the CTD TDP-43, we sought to investigate its effects on the aggregation kinetics of Low Complex Domain of TDP-43 protein (LCD, 263-414aa). Recombinant LCD protein was citrullinated by incubation with PAD4 as described above. The LC MS/MS analysis confirmed PAD-mediated citrullination of 4 out of 5 existing arginine: R268, R272, R275 and R293 TDP-43 residues but not R361 (Fig. 4d, Table 1). AlphaFold renderings of CTD TDP-43 (267-414aa, PDB ID:7KWZ) demonstrated spatial orientation and distance at R268, R272 (Fig. 4e, 15Å) and R275, R293 residues (Fig. 4f, 5Å) as well as hydrogen bonds formed with the adjacent amino acid.

Contrary to the amorphous structure for TDP-43 protein, TEM imaging revealed fibrillar amyloid LCD aggregates, as ThT analysis confirmed max aggregation intensity was reached between 24 - 30 hours (Fig. 4g, i). However, citrullination of LCD disrupted β-sheet aggregates and yielded similar granular structures as the full length citR TDP-43, and distinct from the LCD fibrils (Fig. 4h, magnified images). ThT-based aggregation kinetics studies surprisingly revealed no changes in maximum ThT signal intensity between both proteins (% ThT signal intensity, Fig. 4i) and a right shift in the citR LCD aggregation curve. Analysis revealed a significant increase in citR LCD half-life (*t_1/2_*); LCD *t_1/2_* = 16.89 *vs.* citR LCD *t_1/2_* = 40.80 hours (Fig. 4i and j, p < 0.001). Further, the lag phase (*t_lag_*) of LCD vs. citR LCD aggregation curve calculated by the linear intersection of the slope at *t_1/2_* demonstrated a 6-fold increase in the citR LCD lag time compared to the unmodified LCD (Fig. 4k, LCD *t_lag_*=1.80 *vs.* citR LCD *t_lag_*=12.04 hours, p < 0.01).

Together, our findings demonstrate the biophysical impact of citrullination on TDP-43 kinetic dynamics, where we propose citrullination as a key molecular factor in extending the lag phase and generation of a magnitude of high free-energy primary nuclei resulting in densely packed citR TDP-43 condensates.

### Citrullination of TDP-43 LCD facilitates droplet maturation and transition into liquid-solid condensates

Post-translational modifications alter protein net charges and can act as regulatory switches to LLPS dynamics, influencing their physiological functions^90^. Additionally, considering the arginine’s role in stabilizing inter and intra-relationships with nucleic acid (RNA) and sequence amino acids^86,91^ and interpretations from PONDR stimulations (Fig. 4), we sought to investigate the effects of citrullination in liquid – liquid phase separation (LLPS) kinetics *in vitro.* Unmodified and citR LCD proteins were incubated with RNA for 72 hours following LLPS protocols previously reported^49^. Brightfield and ThT fluorescence imaging under the GFP channel (AF488) were collected to study droplet formation and maturation with time (aging). Images revealed aggregation of 40 µM LCD protein 0 – 72 hrs. time points (Fig. 5a), and following incubation with 250 ng/µl RNA. Interestingly, the unmodified LCD-LLPS droplets were dispersed in small LLPS droplet as early as 2 hours and matured into starburst-like aggregates by 24 hours (Fig. 5b). Surprisingly, brightfield and GFP channel images revealed a lack of citR LCD protein assemblies, with initial droplet-like condensates first appearing by 24 hours (Fig. 5a, bottom panel). Aging of LCD droplets to 72 hours displayed morphologically larger and more concentrated condensates (Fig. 5b). In agreement with the TEM results, images showed that citR LCD (40 µM) droplet size and area coverage was unaffected by RNA (Fig. 5a *vs.* 5b, bottom panel). These findings suggested that citrullination impairs protein interactions with RNA at the expense of droplet maturation into a liquid-solid phase separated condensates (LSPS).

**Figure 5.**
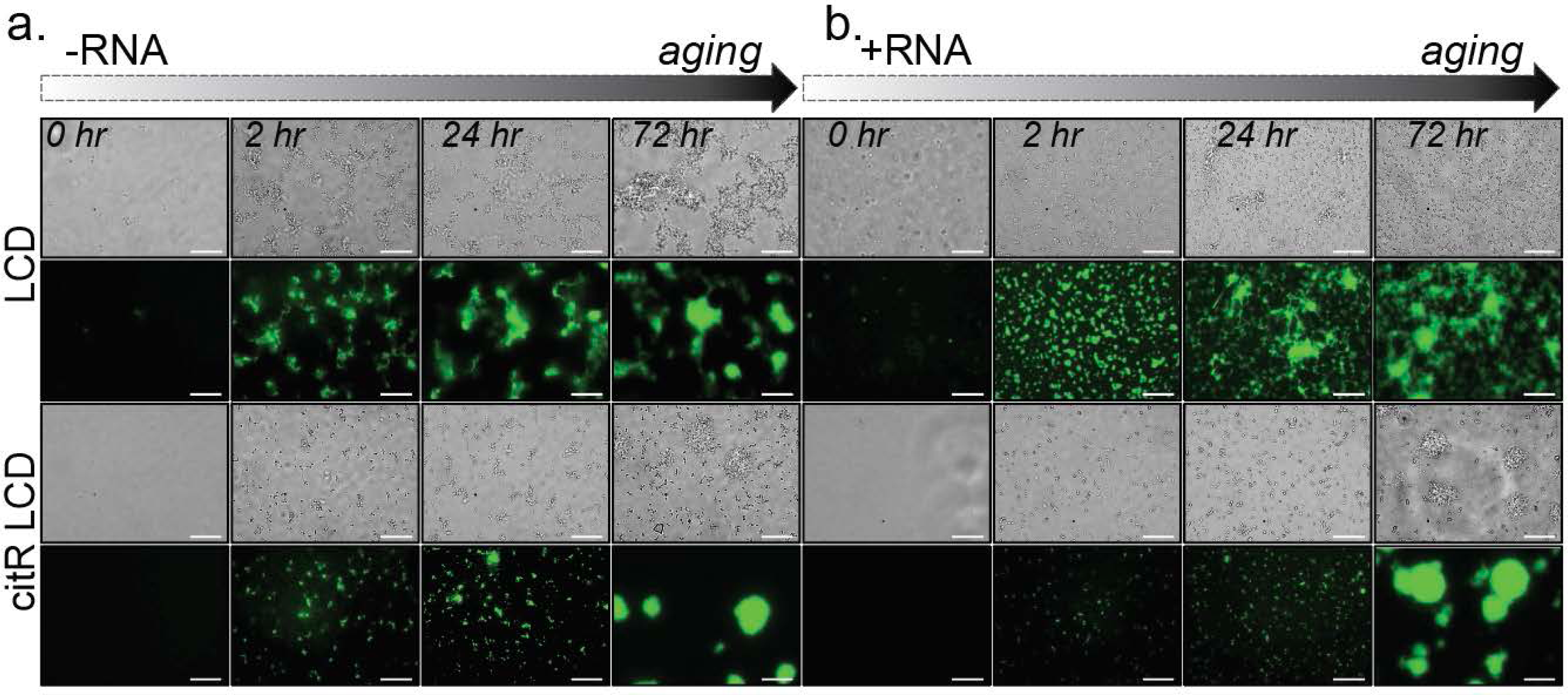
CitR LCD facilitates droplet maturation and aging into liquid-solid phase separated condensates. a, Bright field and GFP fluorescent images of 40 µM LCD or citR LCD incubated between 2 - 72 hours or b, with 250 ng/µL RNA. LCD starburst aggregates formed under RNA – dependent conditions, as citR LCD maturation of condensates was delayed until 72 hours and independent of RNA. n = 3, scale bar is 20 µm.

LLPS interactions are relatively weak and sensitive to changes in temperature and ionic strength, PTMs and protein concentration^92–94^, and the protein kinetics analysis above suggested induced high-energy state of citR LCD. Therefore, we sought to investigate the long-term effects of citrullination on protein: RNA interactions. We incubated LCD and citR LCD proteins at various protein concentrations (10, 20 and 40 µM) with yeast RNA (0, 31.25, 250 ng/µl) for 72 hours (Fig. 6a). Here, we demonstrated that RNA concentration - dependently facilitated starburst-like irregular LCD droplets (Fig. 5b), consistent with TDP-43 LCD maturation or “aging” shown in prior studies^49,95^. However, citR LCD sample was devoid of interactions with RNA at 10 - 20 µm protein concentration (Fig. 6a). Again, we observed maturation of citR LCD droplets first at 72-hour interval and only at 40 µm citR LCD sample. Images demonstrated citR LCD condensates that morphologically displayed viscous, spherical profile and independent of increasing RNA concentrations, suggesting that LSPS formation was driven by C_max_ citR LCD / self-crowding effects (Fig. 6a). ThT signal intensity of LCD and citR LCD “aged” condensates was measured as area occupancy within the field of view (FOV, Fig 6b, circle represent condensate area) and condensate area (object) following increasing protein (10, 20 and 40 µm, Fig. 6c) and RNA concentration (0, 32, 25 and 250 ng/µl RNA, Fig. 6d). Our analysis revealed significant increase in LLPS droplet occupancy/ FOV during LCD droplet aging (Fig. 6b, pink circles), while the droplet size (area µm^2^) remains largely unchanged (Fig. 6c, p > 0.05). Interestingly, analysis further demonstrated significant increase in citR LCD LSPS occupancy/ FOV (Fig. 6b, green circles), and the large condensate size as function of protein concentration (Fig. 6c, ^####^p < 0.0001).

**Figure 6.**
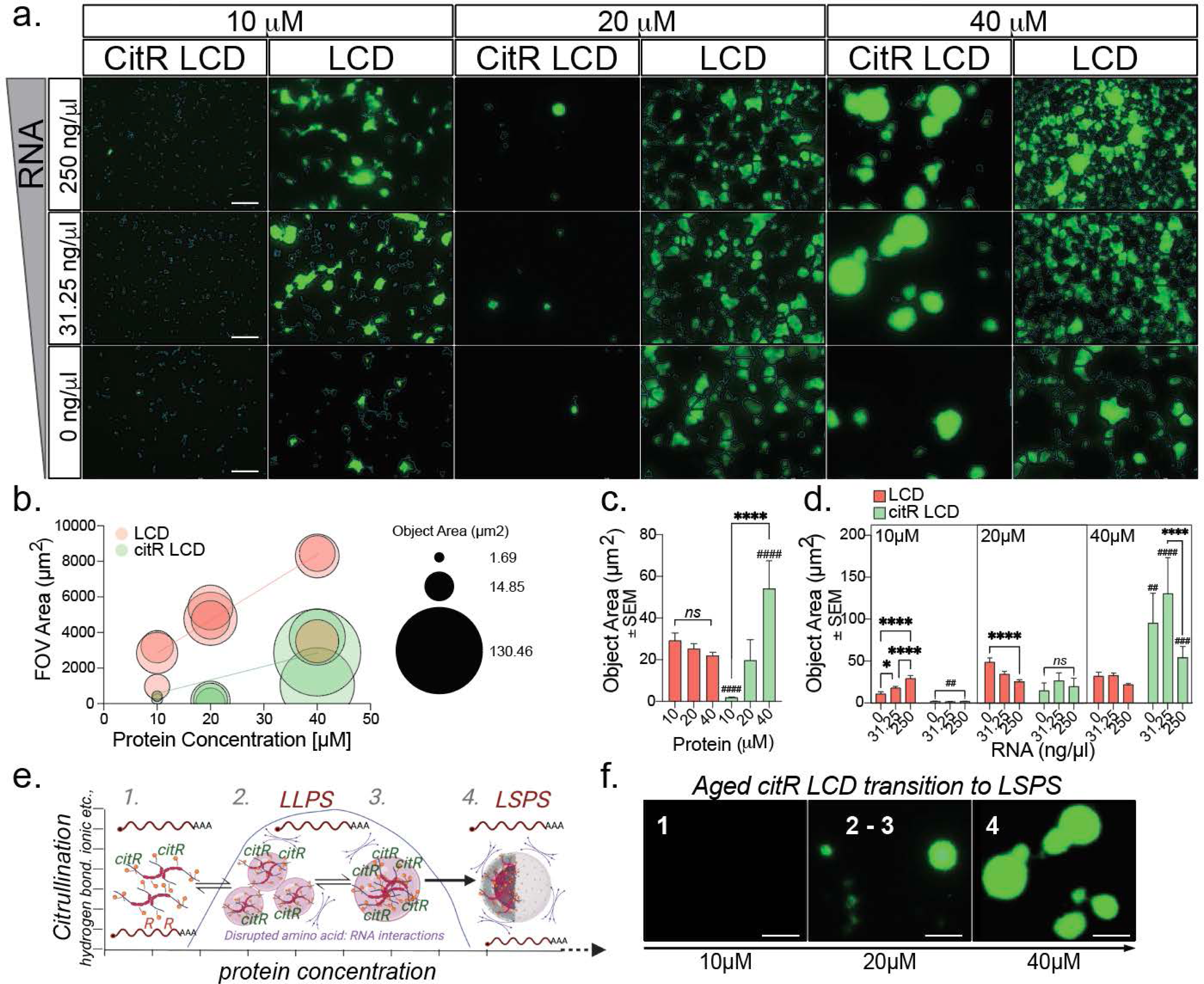
Citrullination of TDP-43 LCD facilitates droplet maturation into liquid-solid phase separated condensates. a, GFP images of LCD or citR LCD aggregated in increasing protein and RNA concentrations at 72-hour endpoint. LCD star-burst aggregate formation with increasing protein and RNA concentration. citR LCD sample florescence signal at the 10, 20 and 40 µM citR LCD protein concentration, and following incubation with increased 0, 31.25 and 250 ng/µl RNA. b, FOV intensity coverage (y-axis) and droplet size for LCD (red circle) and citR LCD (green circle) as function of protein concentration (x-axis). Data represent the linear regression of samples incubated with RNA (250 ng/µl); circle diameter indicates mean object size. c, Mean droplet size for 10, 20, and 40 µM LCD / citR LCD in the presence of 250 ng/µl RNA at 72 hours. Data represent the mean SEM, one-way ANOVA, followed by Tukey’s *post hoc* multiple comparisons test, n = 3, ****p < 0.0001 within the group; #### p < 0.0001 across different groups. d, Mean object size of droplets formed in the presence of increasing concentrations of RNA at 10, 20, and 40 µM LCD and citR LCD at 72 hours. Data represent the mean SEM, two-way ANOVA, followed by Tukey’s *post hoc* multiple comparisons test, n = 3, *p < 0.05, ****p < 0.0001 within same group; ## p < 0.01, ### p < 0.001, #### p < 0.0001 across different groups. e, Proposed model of LLPS- LSPS transition of citR TDP-43 under the two-phase system model. f, Representation of citR LCD transitions with time and protein concentration from our experiment. Scale bar in all images is 20 µm.

Next, we analyzed the LCD vs. citR LCD “aged” condensate size (object area) as a function to RNA concentration. Analysis demonstrated that LCD transition to LLPS was dependent on RNA concentration (Fig. 6d, ****p < 0.0001), while these interactions weakened as LCD protein concentration increased (Fig. 6d, 20 µm, ****p < 0.001, 40 µm, p > 0. 05). We found an opposite trend in the citR LCD sample, where % citR LCD LSPS was strongly driven by protein concentration and independent of RNA (Fig. 6d, 10 and 20 µm, *p > 0.05, 40 µm, ^###^p < 0.001).

Considering our findings, we proposed a two-phase system model by which citrullination serves as an irreversible molecular switch for TDP-43 LLPS-LSPS transitions *in vitro* (Fig. 6e). We posit that modification of the TDP-43 protein net charge repeals pi-pi interactions with RNA and inhibits inter/intra linear pi-pi interactions with the nearby amino acids, extending the nucleation phase (state 1). This event facilitates transition to an intermediate LLPS droplet (state 2 -> 3). Following droplet maturation phase and high TDP-43 protein concentrations, citR TDP-43/LCD protein reaches C_max_ at which state LLPS droplets undergo maturation - “aging”, resulting in loss of regional elasticity and transition to viscous LSPS condensates (state 4). Aging of citR LCD to LSPS condensate states found in our study are compiled in Fig. 6f. Overall, the data presented here suggests the mechanism of how citrullination, by the virtue of reducing protein’s- net charge and self-repealing modifies the concentration of the reactive species and their stacking into solid condensates.

### Generation and validation of citR TDP-43 polyclonal antibodies

The lack of site-specific citrullinated antibodies against a wide range of proteins has limited the ability to investigate the biological effects of citrullination during physiology and disease biology. Considering our findings on the impact of TDP-43 citrullination on protein structure and kinetic, we rationed that a targeted strategy of site-specific citR TDP-43 antibodies will expand our ability to fully investigate epitope and domain-specific cellular mechanisms. Thus, we generated polyclonal citR TDP-43 antibodies against 7 of 11 identified citR epitopes positioned within each functional TDP-43 domain – citR83 (NLS), citR165 and citR191 (RRM1, RRM2), citR268/272, citR275, citR293 (CTD). Details of each citR TDP-43 epitope sequence, localization and domain function are shown in Fig. 7a and Table 3. Antibody specificity to citR epitopes was determined biochemically using similar stoichiometry between recombinant TDP-43: PAD4
 proteins described above. Western blot images revealed the specificity of citR TDP-43 antibodies to the citrullinated TDP-43, as shown by the presence of the 43 kDa band (TDP-43 + PAD4 sample) compared to the unmodified TDP-43 protein. Notably, citR83 (NLS) and citR268/272 (CTD) antibodies recognized the 130 -170 kDa high molecular weight bands (HMW) (Fig. 7b). We found that citR165 (RRM1) antibody recognized the multimer 72 kDa band but bound weakly to the 43 kDa citR TDP-43 monomer. Interestingly, the citR191 antibody (RRM2) demonstrated a reverse preference for the 43 kDa citR TDP-43 species compared to the 72 kDa multimer (Fig. 7b).

**Figure 7.**
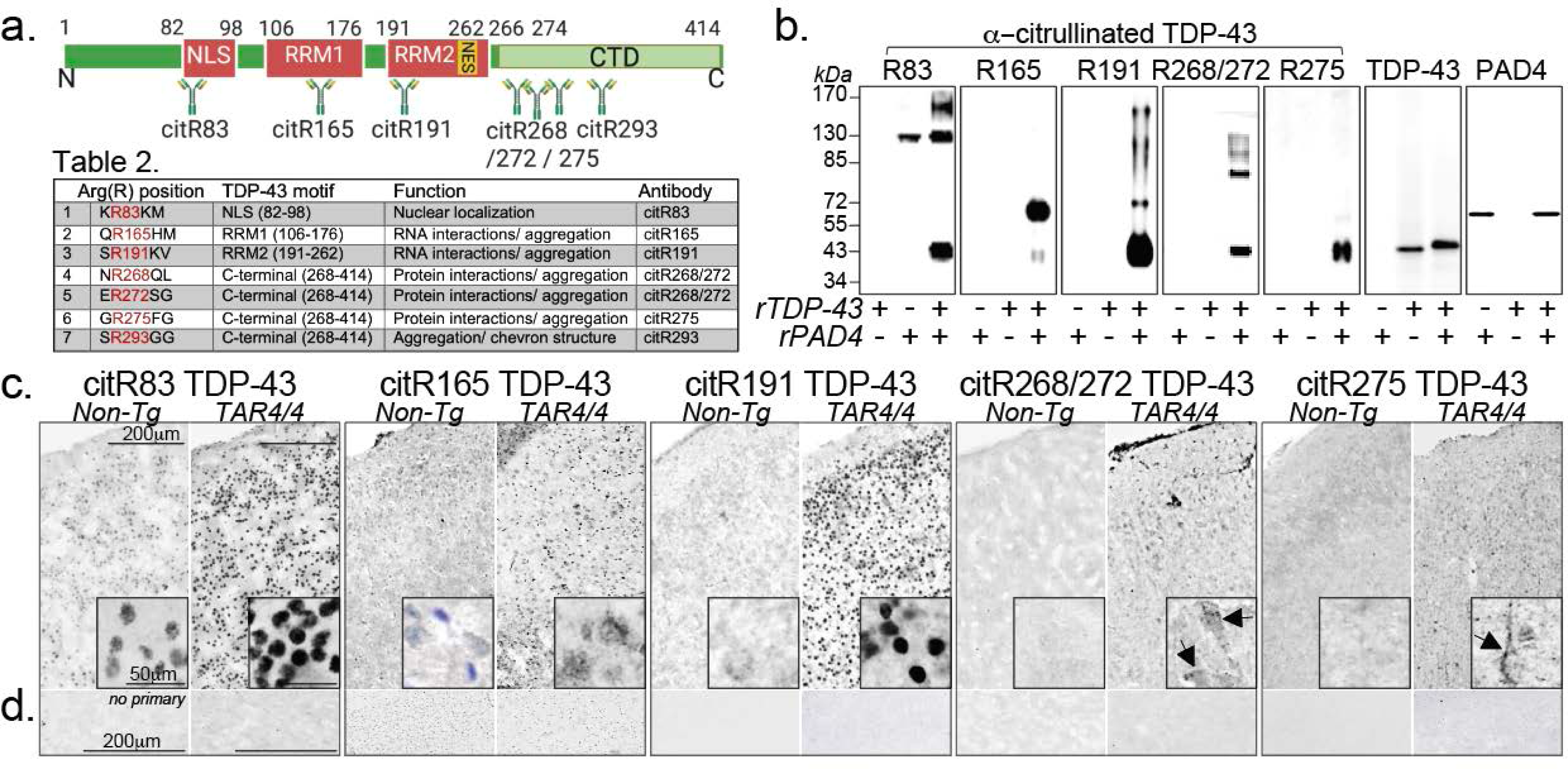
Citrullinated TDP-43 antibody development and validation in vitro and in vivo systems. a, Schematic representation of seven polyclonal antibodies raised against the citrullinated arginine epitopes. Epitope positions within each TDP-43 functional domain is indicated. b, Antibody specificity was determined by western blot via PAD4-mediated citrullinated TDP-43 protein probed with citR83 (NLS), citR165 and citR191 (RRM1, RRM2), citR268/272, citR275 (CTD) antibodies. c, Immunohistochemical images of TAR4/4 and non-Tg littermates cortical tissue labeled against citR TDP-43 antibody panel. Increased neuronal expression of citR83, citR165, citR191, citR268/272 and citR275 in TAR4/4 tissue compared to Non-Tg tissue. Arrows indicate the citR TDP-43 morphologies observed with the citR268/272 and citR275 antibodies raised against TDP-43 C-terminal domain (arrows). Insets represent magnified neuronal citR TDP-43 morphological properties recognized from each antibody. d, Images of cortical tissue incubated with secondary antibody only - “no-primary” control. n = 3, scale bar is 50µm and 200 µm.

Next, we validated the citR TDP-43 antibody specificity via immunohistochemistry labeling of brain tissue from heterozygous TAR4 (∼1.2-fold human) and homozygous TAR4/4 (∼2-fold human gene expression) mouse models and age matched non-transgenic littermates^57^. Overall, the citR TDP-43 antibody panel recognized epitope-specific citrullination of TDP-43 in gene-dose dependent fashion in TAR mice compared to the Non-Tg littermates, while “no primary” control tissue lacked immunoreactivity (Fig. 7c and Supplementary Figure 3). Histochemical images further demonstrated strong positive neuronal signal for citR83 and citR191 antibodies in the cortex of TAR4/4 mice. Interestingly, citR TDP-43 antibodies raised against R268/272 and R275 epitopes within CTD recognized a distinct granular pattern of citR TDP-43 within the neuron (arrows). It is intriguing to suggest that citR TDP-43 antibodies within the intrinsic disorder domain (IRD) may recognized unique structural citR TDP-43 condensates harboring citrullination, similar to our in vitro findings (Fig. 4-6). Collectively, our data demonstrate that citR TDP-43 specific antibodies are useful tools to investigate TDP-43 citrullination at the mechanistic level and link specific citR TDP-43 conformers to pathology *in vivo*.

### Increased citrullinated TDP-43 signatures in the animal models of TDP-43 proteinopathy

To further investigate the citR TDP-43 signatures following TDP-43 pathological progression we performed a detailed immunohistochemical characterization of citR83 (NLS), citR191 (RRM2) and citR268/272 (CTD) TDP-43 levels in the TAR mouse model. Cortical and hippocampal images of the cortical layers 4-5 (Fig. 8b, d, f, and insets) and hippocampal CA1 layer (Fig. 8i, k, m, and insets) demonstrated increased neuronal expression for citR83, 191 and 268/272 TDP-43 levels in TAR4/4 mice compared to TAR4 and non-transgenic control mice. High magnification images further revealed epitope-specific morphological characteristic for each citR TDP-43 species *in vivo*; citR83 TDP-43 antibody recognizes distinct nuclear and cytoplasmic neuronal TDP-43 inclusions, while citR191 TDP-43 antibody recognized mostly perinuclear TDP-43 conformers; these patterns remained consistent for both brain regions in TAR4/4 mice. Similar to our validation studies, we observed a unique immunoreactivity pattern for the citR268/272 TDP-43 antibody. High magnification images from both CX and CA1 regions revealed granular citR268/272 positive TDP-43 conformers in TAR4/4 mice (Fig. 8e, f, I, m, arrows, and insets). Overall, we found significant gene-dose dependent increase in citR TDP-43 immunoreactivity compared to Non-Tg littermates (TAR4/4 > TAR4, Fig. 8g, n). Analysis of citR83 and citR268/272 TDP-43 % positive area in CX demonstrated over 2.5-fold increase in TAR4/4 mice (Fig. 8g, **p < 0.01, ***p < 0.001, ****p < 0.0001), while citR191 TDP-43 % positive area was increased by 14-fold in TAR4/4 mice compared to the Non-Tg controls (Fig. 8g, ***p < 0.001, ****p < 0.0001). Hippocampal immunoreactivity for citR TDP-43 followed similar trends, as % positive area analysis demonstrated a 2-fold increase in citR83 and citR268/272 levels in the hippocampus of TAR4/4 mice compared to the Non-Tg littermates (Fig. 8n, 1.9 - 2.5-fold, respective, *p < 0.05, **p < 0.01, ***p < 0.001). Similarly, we found that citR191 TDP-43 % positive area was increased by 5-fold in TAR4/4 mice compared to the Non-Tg controls (Fig. 8n, 4.9 - fold change, ****p < 0.0001). Overall, we provided evidence on increased citR TDP-43 profile in TAR model of TDP- 43 proteinopathy and determined epitope-specific effects of citrullination. We propose citR83 residue as a potential PAD4 “restricted epitope” as its levels were increased early in TAR mouse model. Similarly, we identify citR191 TDP-43 as highly responsive to PADs activity and demonstrated the presence of morphologically novel and granular citR268/272 conformers *in vivo*.

**Figure 8.**
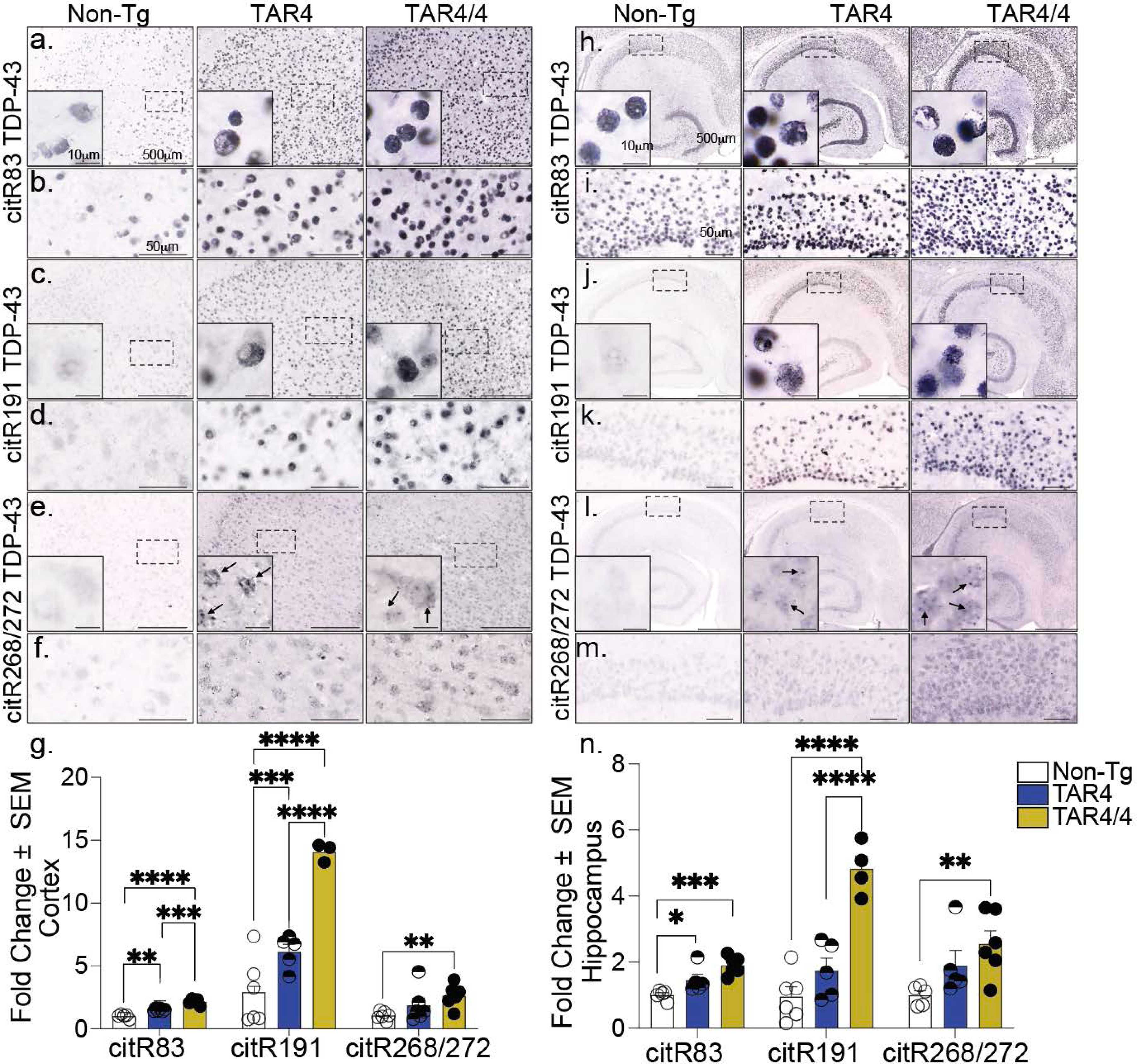
Induced TDP-43 citrullination in TAR mouse model. a-m, Immunohistochemical images of TAR4 and TAR4/4 cortex and hippocampal tissue labeled with a, b, h, i, citR83, c, d, j, k, citR191, and e, f, l, m, citR268/272 TDP-43 antibodies. g, n, Fold change of TDP-43 citrullination levels in the cortex and hippocampus of TAR mice normalized to the Non-Tg control. Data represent the mean of the values ± SEM; Two-way ANOVA, followed by Tukey’s *post hoc* multiple comparisons tests, n = 5, *p < 0.05, **p < 0.01, ***p < 0.001, ****p < 0.0001.

### Increased PAD2 and 4 expression and TDP-43 citrullination profile in TAR mouse model

Next, we asked whether the increased citR TDP-43 profile observed in this model was due to cell-specific PAD expression^32–35^. Utilizing immunohistochemistry labeling we demonstrated increased PAD2 and PAD4 immunoreactivity in the respective astrocytic and neuronal population in the cortex of TAR4/4 mice (Fig. 9a). Analysis of % positive area revealed significant increase in astrocytic PAD2 levels in TAR4/4, but not in TAR4 mice, compared to the Non-Tg littermates (Fig. 9b, 1.9 - fold change, *p < 0.05). We also found significant increase in neuronal PAD4 levels in TAR4/4 mice compared to TAR4 and Non-Tg littermates (Fig. 9b, 6.6 - fold change, **p < 0.01, ****p < 0.0001). Notably, PAD4 expression was significantly higher than the PAD2 in TAR4/4 mice (6.69 - fold change, ***p < 0.001).

**Figure 9.**
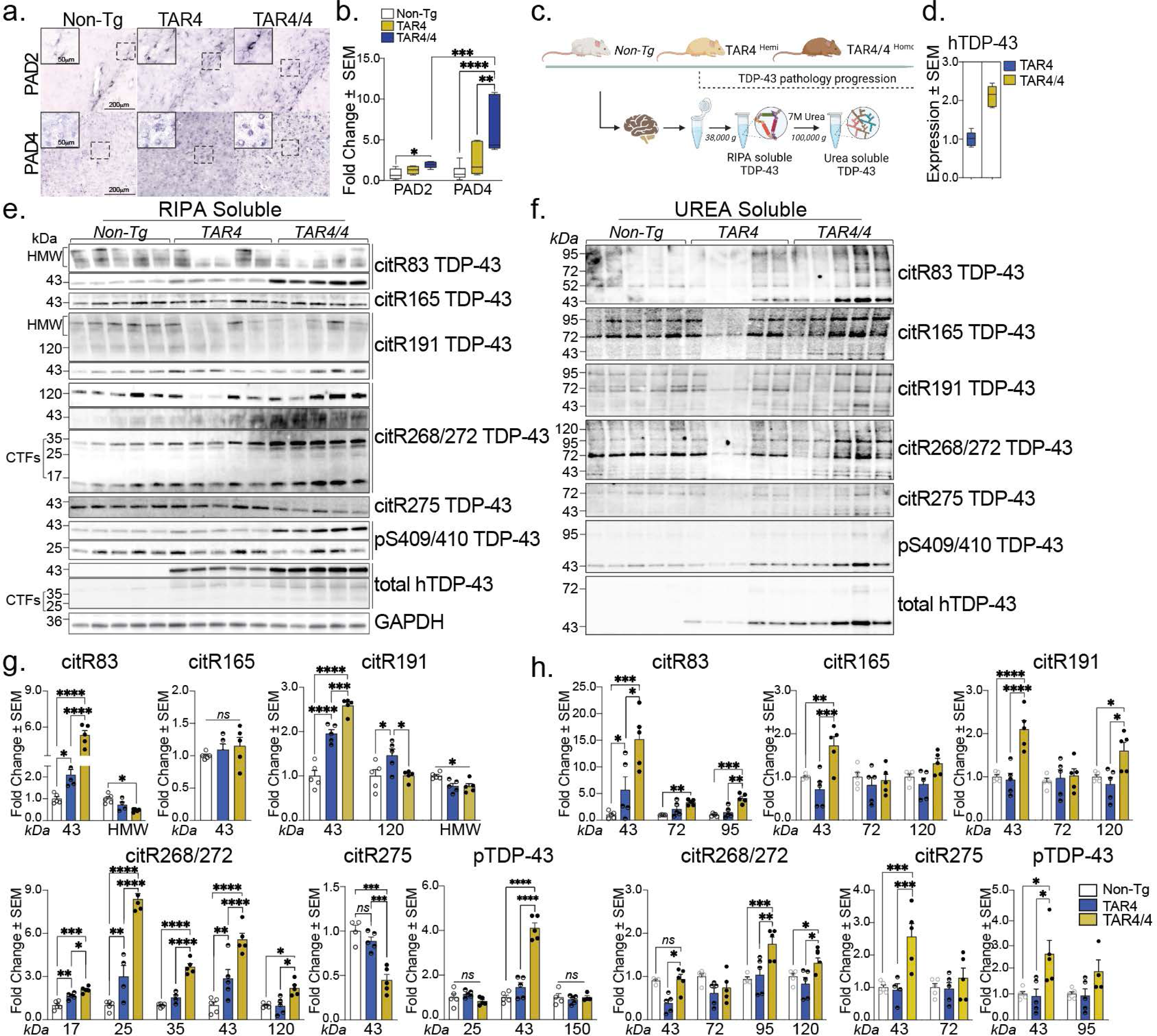
Epitope-specific TDP-43 citrullination and conformer TDP-43 species solubility in TAR mouse model. a, Immunohistochemical images of Non-Tg, TAR4 and TAR4/4 cortex labeled with PAD2 and PAD4 antibodies. b, Fold change of PAD2 and PAD4 (% area mean of the values ± SEM) normalized to the Non-Tg control. One-way or Two-way ANOVA, followed by Tukey’s *post hoc* multiple comparisons tests, n = 5, *p < 0.05, **p < 0.01, ***p < 0.001, ****p < 0.0001. c, Schematic overview of the TAR TDP-43 mouse model and the fractionation of cortical tissue into RIPA-soluble and (7 M) Urea-soluble fractions. d, The expression levels of the human TDP-43 in the TAR model. e, Cortical RIPA soluble and f, Urea soluble fraction analyzed by Western blotting and probed for citR TDP-43 antibody panel (citR83, 165, 191, 268/272, 275), pTDP-43 409/410 and total human TDP-43 protein. g, Quantification of citR TDP-43 43 kDa protein levels, proteolytic fragments (17, 25 & 35 kDa) and intermediate high molecular species (72, 98 & 120k Da) normalized to GAPDH in RIPA and h, Urea fraction normalized to the total protein loaded signal. Data represent the mean of the values ± SEM; One-way ANOVA, followed by Tukey’s or Šidák *post hoc* multiple comparisons tests, n = 5, *p < 0.05, **p < 0.01, ***p < 0.001, ****p < 0.0001.

**Figure 10.**
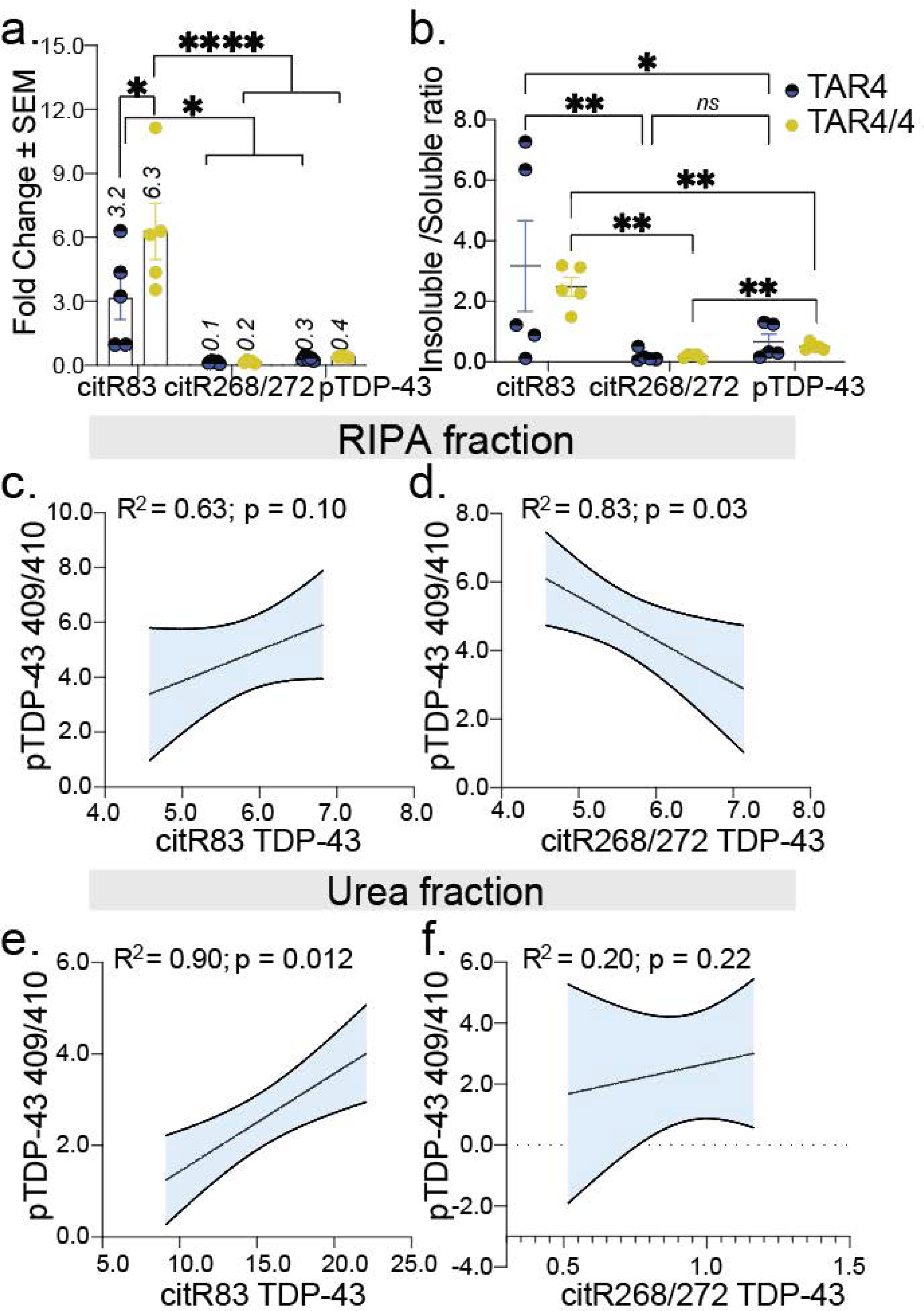
Citrullination drives site-specific TDP-43 solubility *in vivo*. a, Quantification of the 43 kDa TDP-43 species as insoluble to soluble ratio and b, fold change for citR83, citR 268/272 epitopes and pTDP-43 levels in TAR4/4 mice. Values are normalized to Non-Tg littermates and calculated as the mean ± SEM; One-way ANOVA, followed by Tukey’s or Šidák *post hoc* multiple comparisons test, n = 5, *p < 0.05, **p < 0.01, ****p < 0.0001. c, Correlation coefficient for citR83 epitope and pTDP-43 in the RIPA and d, Urea-soluble fractions. e, Correlation coefficient for citR268/272 epitopes and pTDP-43 in the RIPA and f, Urea-soluble fractions. Pearson correlation coefficient (R^2^) was determined followed by Student t-test, p two-tail, linear regression lines with 95% Cl are shown (shaded areas).

To determine whether citrullination regulates TDP-43 protein unfolding and solubility, we performed biochemical RIPA soluble and 7M Urea-soluble fractions fractionation assays as we previously reported^59^ (Fig. 9c). Western blot analysis from cortices of TAR4, TAR4/4 mice and age-matched non-transgenic littermates confirmed the total human TDP-43 expression in TAR4/4 mice (Fig. 9d, ∼2 - fold gene expression hTDP-43, p < 0.001)^57^. Both RIPA and Urea fractions were then probed with epitope-specific citR TDP-43 antibody panel: citR83, citR165, citR191, citR268/272, and citR275 (Fig. 9e, f). We included phosphorylated S409/410 TDP-43 (pTDP-43) as a pan marker of pathology^2^. The analysis of RIPA-soluble fraction demonstrated a significant increase in citR TDP-43 levels gene-dose dependent (TAR4/4 > TAR4) and compared to the age-matched Non-Tg littermates (Fig. 9g). Analysis of different molecular weight TDP-43 conformers demonstrated a significant increase in the 43kDa citR83 TDP-43 levels in TAR4 and TAR4/4 mice (Fig. 9e, g, 2 - *vs.* 5 - fold change, **p < 0.01, ****p < 0.0001, respectively), citR191 (1.9 - *vs.* 2.5 - fold change, ****p < 0.0001), and citR268/272 (2.8 - vs. 6 - fold change, *p < 0.05, ****p < 0.0001) compared to the Non-Tg littermates. We found no changes in the levels of citR165 TDP-43, while a significant reduction in citR275 TDP-43 levels was measured in the soluble fraction of TAR4/4 mice compared to the Non-Tg littermates (Fig. 9e, g, ***p < 0.001). As expected, we found increased pS409/410 TDP-43 levels (43 kDa band) in TAR4/4 but not TAR4 mice (Fig. 9e, g, ****p < 0.0001 *vs.* p > 0.05, respectively). Interestingly, citR268/272 TDP-43 antibody recognized several C-terminal proteolytic species; 17 kDa, 25 kDa, and 35 kDa bands/ CTFs, which were significantly increased in TAR4/4 mice (Fig. 9e, g, TAR4/4 > TAR4 mice, *p < 0.05, **p < 0.01, ***p < 0.001, ***p < 0.0001). The 25 kDa CTF species were modestly recognized by the pS409/410 TDP-43 antibody in TAR4/4 mice but did not reached significance. Meanwhile, analysis of the high molecular weight species; 95, 120 and 130-170 kDa bands (HMW), demonstrated significant reduction in HMW citR83 and citR191 TDP-43 levels (*p < 0.05), while significant increase was found only for the 120 kDa citR268/272 conformer in the soluble fraction of TAR4/4 compared to TAR4 and Non-Tg littermates (Fig. 9e, g, *p < 0.05).

Next, we analyzed citR TDP-43 abundance in the urea-soluble fractions from TAR mice cortices as an indicator of citR TDP-43 aggregation *in vivo*. Surprisingly, we found a significant increase in 43 kDa citR83 TDP-43 levels (Fig. 9f, h, 15-fold, *p < 0.05, ***p < 0.001), as well as 72 and 95 kDa species (3 - 4-fold change, respectively, ***p < 0.001, **p < 0.01) in TAR4/4 mice but not TAR4 and Non-Tg littermates. Further, we found significant accumulation of 43 kDa TDP- 43 citR165 (1.7 - fold, *p < 0.05), citR191 (2 - fold, ***p < 0.0001), and citR275 TDP-43 in TAR4/4 mice (2.5 - fold, ***p < 0.001), while 43 kDa TDP-43 citR 286/272 levels remained unchanged compared to the Non-Tg control. Instead, a significant accumulation of HMW conformers; 95 kDa and 120 kDa citR268/272 TDP-43 (1.8 -fold, *p < 0.05) and 120 kDa citR191 TDP-43 (1.6-fold, *p < 0.05) was measured in TAR4/4 mice in this fraction (Fig. 9f, h). To note, we found accumulation of the 43 kDa insoluble pS409/410 TDP-43 but not HMW (95kDa band) in TAR4/4 mice compared to TAR4 and Non-Tg control (Fig. 9f, h, *p < 0.05).

To determine the effects of epitope-specific citrullination on TDP-43 solubility index in this model, we analyzed the ratio of insoluble to soluble for citR83 and 268/272 TDP-43 (43 kDa species) and compared to that of pTDP-43 canonical hallmark. Surprisingly, we identified citR83 TDP-43 as the dominant epitope driving TDP-43 insolubility *in vivo* (Fig. 10a, mean 2.5 ± 0.3, **p < 0.01) compared to citR 268/272 (mean 0.2 ± 0.03) and pTDP-43 (mean 0.5 ± 0.05) in TAR4/4 mice. In fact, when normalized to the Non-Tg levels we found 3.2 *vs.* 6.3-fold increase in citR83 TDP-43 (43 kDa) levels in TAR4 and TAR4/4 mice, respectively, compared to 0.3 *vs*. 0.4-fold increase for pTDP-43, respectively (Fig. 10a, *p < 0.05, ****p < 0.0001). Citrullination of the R268/272 epitope resulted primarily in accumulation of soluble species (Fig. 10b. 0.1 v*s.* 0.2-fold, respectively, *p < 0.05, **p < 0.01). Correlation of pTDP43 to citR83 TDP-43 levels revealed a positive correlation for RIPA (Fig. 10c. R^2^=0.63, p_two-tailed_ = 0.1045) and a stronger positive correlation for urea fractions (Fig. 10e. R^2^=0.99, p_two-tailed_ = 0.0128) between the two PTMs. Not surprisingly, we found a strong negative correlation between pTDP-43 and citR268/272 in the RIPA fraction (Fig. 10d, R^2^=0.83, p_two-tailed_ = 0.0304), while no correlations were found in the urea fraction (Fig. 10f, R^2^=0.20, p_two-tailed_ = 0.2244).

Overall, our data indicated abundant citR TDP-43 levels *in vivo* following the pathology progression of TAR mouse model. We identified citR83 - induced precipitation of citR TDP-43 insolubility *in vivo*. Both IHC and biochemical analysis indicated that citR 268/272 in the LCD drives formation of soluble, granular puncta proteoforms, strongly suggesting epitope-dependent effects of citrullination in this model.

### Defining the citR TDP-43 signatures in LATE- NC and ADNC+LATE co-pathologies

We next investigated the citR TDP-43 immunohistochemical results in comparison to TDP-43 pathology in LATE-NC (with or without comorbid ADNC) in human brain sections^6^. We performed immunohistochemical labeling and imaging of brain sections from healthy control cases (HC) and those presenting ADNC, LATE - NC stages 1, 2, or ADNC + LATE - NC.

Demographic information and neuropathological diagnoses of the selected tissue from 19 research participants are depicted in Supplementary Table 2. Brain images from these cases revealed robustly increased immunoreactivity for citR83, citR191 and citR268/272 TDP-43 in LATE-NC and ADNC + LATE-NC cases, with no/low immunoreactivity observed in in pure ADNC or HC cases (Fig. 11a). The citR TDP-43 immunoreactivity was quantified as percent area intensity within seven regions of interest (ROIs) / brain area representing each disease state (LATE-NC stage 1 = amygdala, LATE-NC stage 2 = amygdala + hippocampus)^6^ and compared to pTDP-43 antibody staining (1D3 clone, pS4109/410) (Fig. 11b). The relationships between citR TDP-43 intensities were mapped across disease states, and we identified epitope-specific differences in LATE - NC (stage 1 and 2) and ADNC + LATE - NC tissue compared to AD or HC brain tissue (Fig. 11b, Supplementary Fig. 4, a1-a10, statistical values depicted in Supplement Table 3). For example, we found that citR83 TDP-43 levels were increased in LATE - NC stage 1 and 2, reaching the highest levels in ADNC + LATE - NC cases (Supplementary Fig. 4, a1, a9 - a10, Supplementary Table 2). By contrast, we found that citR191 and citR 268/272 TDP-43 levels were prominent in LATE - NC stage 2 and significantly higher than pTDP-43 levels (Supplementary Fig. 4, a2 – a4, a8), but was not exacerbated in ADNC + LATE-NC individuals (Supplementary Fig. 4, a7-a10 and Supplementary Table 2).

**Figure 11.**
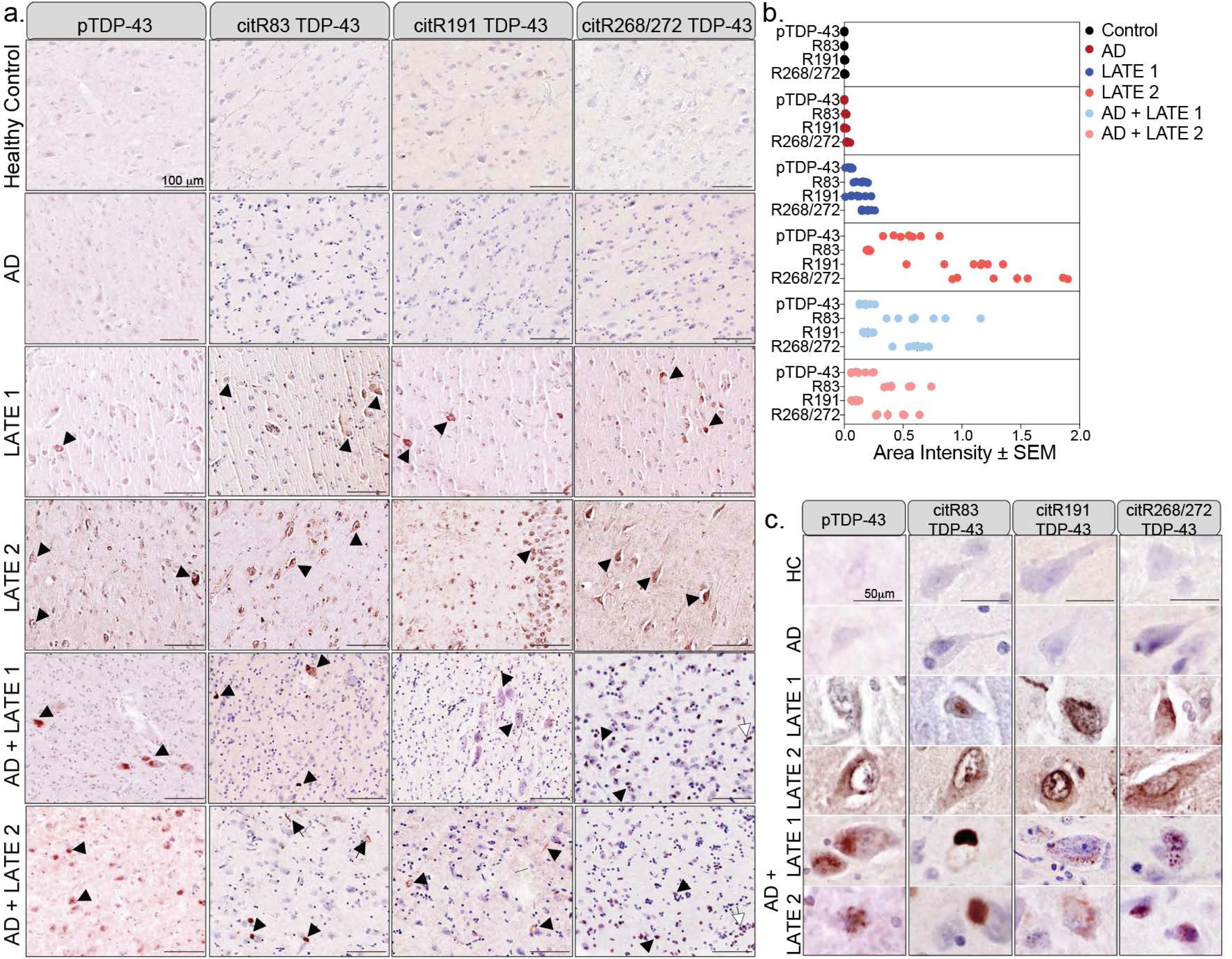
Epitope-specific citrullination of TDP-43 reveals regional and temporal neuropathological changes in LATE and ADNC + LATE human brain. a. Images of brain tissue form healthy controls (HC) and LATE-NC, ADNC + LATE-NC disease cohorts and labeled with pTDP-43 and citR TDP-43 antibody panel. Citrullination signatures of citR83, citR191 and citR268/272 TDP-43 antibodies defined the nuclear and cytoplasmic inclusions found in LATE-NC and ADNC + LATE-NC disease states (arrowhead) while glial labeling was also observed (white arrow). b, ROI (n = 7/ brain regions) intensity ratio to HC of each citR TDP43 species and pTDP-43 in brain regions defined by each disease state. c, Magnified images of representative citR morphologies in each disease state. Scale bars are 100 and 50 µm. *Healthy Control (HC), Alzheimer’s Disease* neuropathologic changes (ADNC)*, Limbic-Predominant Age-Related TDP-43 Encephalopathy* neuropathologic changes (ADNC) *(LATE - NC). LATE stage 1 = amygdala, LATE stage 2 = hippocampus, ADNC and HC = frontal cortex*.

Morphologically, we found citR83 TDP-43 immunoreactivity in neuronal nuclear / perinuclear membrane in early LATE-NC stages, precipitating into large TDP-43 cytoplasmic inclusions (NCIs) in ADNC + LATE-NC cases (Fig. 11c). Interestingly, citR191 TDP-43 was found largely in perinuclear membrane and neuronal layers of dentate gyrus and hilus in LATE-NC stage 2 (Fig. 11b, c). Finally, citR 268/272 TDP-43 immunoreactive structures were largely represented as granular puncta in LATE-NC cases (Fig. 11b, c), resembling similar immunoreactivity profile obtained in the TAR model (Fig. 8).

While the pathogenicity of newly identified citR TDP-43 epitopes remains unknown, we utilized the AlphaMissense algorithm in AlphaFold2 to predict the likelihood of each citR epitope to render pathogenicity in human disease^96^. In this algorithm, we treated each targeted R → Q epitope as a “missense mutation”, predicting a pathogenetic score for R83 → Q “mutation” within the NLS domain of 0.998. When compared to other potential missense mutations of the R83 position, we found a high residue pathogenicity score that ranged between 0.98_min_ – 1.00_max_ (Supplementary Fig. 6a). The pathogenicity scores for the RRM epitopes were also predicted to be relatively high; R165 → Q “mutation” score = 0.978 (Supplementary Fig. 6b, overall residue pathogenicity = 0.917_min_ – 1.00_max_), R191 → Q “mutation” pathogenetic score = 0.737 (Supplementary Fig. 6c, residue pathogenicity = 0.737min - 0.998max). Surprisingly, R268, R272, R275 and R293 → Q “mutations” within CTD did not yield a pathogenetic score (Supplementary Fig. 6d, respective residue pathogenicity = 0.626 - 0.860 *vs.* 0.561 - 0.838 *vs.* 0.643 - 0.885 *vs.* 0.643 - 0.894).

Overall, these findings not only demonstrated citrullination of TDP-43 along the pathology progression in human diseases but also open new venues to investigate citR epitope and regional propensity to pathogenicity in our future studies.

## Discussion

We identified citrullination of TDP-43 as an apparently irreversible post-translational modification associated with LATE–NC pathogenesis in human brains and mouse models. We presented evidence for the PAD4 and PAD2 - mediated citrullination in 1) human TDP-43 in NLS, RRMs, and LCD functional domains, and demonstrated its role on 2) unfolding of TDP-43 structure, 3) protein lag phase dynamics and assembly kinetics along with mechanistic insights on the effects of CitR on 4) TDP-43 LLPS droplet aging and the transition to LSPS condensates. Additionally, we showed increased PAD2 and PAD4 expression in animal models following TDP-43 pathology progression, 5) developed and validated several epitope-specific citR TDP-43 antibodies, which were key to determining 6) epitope and domain-specific role of citrullination on TDP-43 protein solubility and pathogenicity *in vivo*. Importantly, we provided key evidence on 7) TDP-43 citrullination and its signatures during TDP-43 proteinopathy in pure LATE-NC and ADNC + LATE-NC. Considering these findings, we propose that the irreversible nature of citrullination contributes to a new and unique form of TDP-43 pathology. Notably, we demonstrate here that irreversible effects of citrullination are largely epitope and domain – dependent and discover at least two citR TDP-43 proteoforms; citR83 and citR268/272 with disparate but consequential effects on TDP-43 proteinopathy.

Advances in mass spectrometry technology have enabled detailed analysis and identification of most commonly reversible PTMs; serine phosphorylation, lysine acetylation and ubiquitination, proteolytic cleavage fragments were isolated from TDP-43 insoluble fractions from ALS and FTLD-TDP brain tissue^2,3,97–101^. Targeted LC MS/MS analysis have been crucial to identifying and mapping the PTM profile of less studied TDP-43 reversible modifications, such as asparagine/ glutamine deamidation and methionine oxidation, in ALS patients^97^. In contrast, citrullinated proteins are generally present in relatively low abundance and their signals are largely suppressed by other molecules^102^. However, several methodological advancements that included identification of HNCO moiety elimination in b- and y-fragmented ions as a marker, led to high confidence discovery of citrullination sites and citrullinome profiling in multiple tissue^102–105^. These data-driven strategies were necessary for propelling the citrullinome studies in cell biology; from creation of reference spectra library to mining of human tissue and discovery of over 300 new citrullination sites from human proteins^106^. Similar successes recently led to citrullinome profiling in PAD2 and PAD4 overexpressing models^107^ and mining of the citrullinome in HL60 leukemia cells identifying 14,056 unique citrullination sites in over 4000 proteins^69^.

Considering the large interest in targeting PAD enzymes in a variety of diseases^108,109^, we demonstrated here increased expression of PAD2 and PAD4 enzymes in TDP-43 mouse models, followed by increased levels of citrullinated TDP-43. Interestingly, recent work from our laboratory demonstrated increased neuronal PAD4 mRNA and protein expression in hippocampal and cortical regions of AD patients^73^, confirming earlier reports^27,35,43^. We further demonstrated that increased PAD4 expression correlated with tau pathology in AD patients proposing that increased PAD4 profile might be implicated with ADNC and other tauopathies. In agreement, *PADi4* SNP rs16824888 is associated with elevated risk for AD in the Caucasian population (*The National Institute on Aging Genetics of Alzheimer’s Disease Data Set)*, while *PADi4* SNP rs2240340 is a risk factor for rheumatoid arthritis^110^. Induced astrocytic PAD2 levels were also reported in the brain tissue from ALS and in AD patients^32,73^. Conversely, one study reported low PADi4 mRNA expression of C-allele SNP rs2240335 (no changes in protein expression), which was associated with increased ALS risk and early disease onset^47^. In the study, PAD4 – mediated citR of FET proteins favored FUS and hnRNPA1 protein solubility and lessened FUS accumulation in SGs in cellular models. The authors identified the consensus RG/RGG motifs in FET proteins were preferred by the PAD4 activity, but TDP-43 was not among the possible targets. In our study we demonstrated citrullination of 11 Arg epitopes across TDP-43 amino acid sequence and found two best-fit RX(XR) and R(X)G/RGGG sequences targeted by PADs with equal intensity (Supplement Table 1). Surprisingly, while RX(XR) motif was prominent across NTD and RRMs protein regions, R(X)G/RGGG motif was only present in the 273-317 aa (Gly/Ser-rich) segment of CTD. These contradictory findings could possibly reflect the difference in PAD’s Ca^2+^ - dependent activation states, different methodologies and/ or substrate specificities (i.e., TDP-43 *vs.* FUS/FET). Within the frame of published findings and developments in PAD - specific drugs tested in cellular and animal models^109,111,112^, we suggest that our study crucially highlight the renaissance of PAD4 (and PAD2) as therapeutic targets across the neurodegenerative disease spectrum.

Here, high mass accuracy and manual examination of the MS spectra provided the first site-specific atlas of TDP-43 citrullination at 55% of R residues within its amino acid sequence. We found similar PAD2 and PAD4 citrullination of epitopes across TDP-43 domains; NTD (R44, R52, R83), RRM1 (R165, R171), the linker segment (R189), RMM2 (R191) and LCD/CTD region. Interestingly, we identified citrullination of the only three charged R268, R272, R275 residues in the N-terminus as well as the R293 residue in the flat sequence of CTD. Notably, we showed a high PAD2 and PAD4 preference for R293 in recombinant protein, which yielded the strongest confidence score (PEP = 7.76e-270, mass error = - 0.74837). Despite the challenges of detecting low abundance citrullinated protein, IP-mass spectrometry identified citR293 TDP-43 in the hippocampal tissue from TAR4/4 mice, as shown by high quality MS spectra b- and y- fragmented ions (Supplement Fig. 1c). Not surprisingly, a recent study confirmed the presence of citR293 chevron fold by cryo-electron microscopy (Cry-Em) retrieved from the brain of three patients diagnosed with FTLD-TDP type A^113^. Interestingly, citR293 was proposed to be the distinguishing modification for all three FTLD-TDP type A cases from the double-spiral fold of ALS and FTLD-TDP type B. The citrullination of R272 and R275 was not identified, their position was reported in the solvent-exposed and less-ordered segment of the chevron fold. We presented here the additional identification of citR268, R272 and R275 residues and demonstrated their role in formation of distinct LSPS condensates that are uniquely presented as granular puncta in TDP-43 mouse and human brain tissue. Further, citrullination of NLS and RRM motif, demonstrated here by citR83 and citR191 antibodies, displayed largely nuclear and perinuclear immune intensities in the hippocampal and dentate gyros cellular layers of LATE stage 1 and stage 2 individuals. Considering the nuclear presence of PAD4 (and PAD2)^27,114^, our data are the first to suggest citrullination of TDP-43 in the nucleus and nuclear envelope of neurons in early stages of pathology. It is therefore intriguing to suggest that the identification of ciR-atlas combined with the discovery of citR TDP-43 signatures and morphologies might indeed reveal a novel form of TDP-43 proteinopathy associated with human diseases.

To this point, our TEM analysis and resolved 3D tomography revealed the biophysical properties of citR TDP-43 and citR LCD conformers/assemblies *in vitro*, signifying unique formation patterns compared to the unmodified TDP-43 aggregates. The ThT analysis of citR LCD and LCD droplet under aging conditions^49^ revealed extended protein lag-time and intensified liquid-solid condensates independent of interactions with RNA. Not surprisingly, LCD is a highly intrinsic disordered segment of TDP-43 readily undergoing transitions to LLPS^76,115^. However, determining whether these principles differ from those of the amyloid deposition can help provide mechanistic explanation to how citrullination expands the TDP-43 structural diversity and potential pathogenicity. Following the classical amyloidal aggregation curve models, we showed that citR significantly increased citR LCD *t_1/2_*(by 40%) and *t_lag_* (by 80%), suggesting extended nucleation and seeding phase, defined by literature as ∼5% amyloid-like growth^116^. Although, the idea of IDR-initiated amyloid-like structures is consistent with our findings, we identified citR as an essential PTM, which facilitated the transition of otherwise a dynamic LLPS droplet^82^ to highly viscous, liquid - solid condensates. This is consistent with reports demonstrating that the steric interactions of arginine residues govern TDP-43 phase-separation kinetics^117–121^, strongly suggesting that molecular determinants such as citrullination hinders the buffering role of RNA in TDP-43 phase separation in cells and aggregation^82,122,123^. Considering our findings, we propose a hypothetical model suggesting that citrullination and saturation of cellular C_max_ [citR TDP-43], shifts protein dynamics/ homeostasis in favor of aging and formation of irreversible LSPS condensates. Applying biophysical two-phase systems models, we suggest that modification of the positive arginine to neutral citrulline repeals pi-pi interactions with regional nucleic acid and disrupts the inter – and intra linear pi-pi interactions with the primary amino acid such as arginine^86^.

While we did not test citR TDP-43 toxicity in this study, evidence has demonstrated how transition to phase-separated states for intrinsic disorder proteins (incl. TDP43, TAF15, tau, FUS etc.) challenges protein homeostasis and initiate pathological cellular processes^82,124,125^. Prediction algorithms from PONDR/ FuzDrop R → Q “pseudo-citrullination” and epitope/ region-specific AlphaMissense analysis (AlphaFold2) predicted citR-induced “hot -spots” and increased TDP-43 LCD disordered state (p_LLPS_ score = 0.8981 → 0.9456), but low pathogenicity. Interestingly, these predictions conceptually linked the high predicted pathogenicity of citR83 residue with the citR83 TDP-43 NCI morphologies identified in both human and mouse tissue.

Similarly, the predicted low pathogenicity for citR268/272 and citR275 TDP-43 could indeed reflect the gap in knowledge regarding citR LCD LSPSs, as an unknown TDP-43 proteoform.

Leveraging the citR TDP-43 specific antibodies we signified the epitope - specific citR TDP-43 profile in animal models with varying degrees of TDP-43 pathology. Indeed, we identified citrullination of R83 in the minor NLS pocket as a driving PTM to TDP-43 insolubility, and linked the citR TDP-43 morphological profiles with the distinct insolubility *in vivo*. Our data provided a hypothetical mechanistic framework to describe the potential pathogenicity of citR83 in TDP-43 pathology. Considering that R83 resides within the K82R83K84 motif of the minor pocket of nuclear localization signal (mNLS)^126,127^, its steric interactions with importin α1 are proposed to be mechanistically necessary^128^. Indeed, mutations or PTMs in proximity to R83 destabilized binding to importin α1 and overall reduced the NLS backbone dynamics, suggesting the biological role of R83 in TDP-43 nucleocytoplasmic shuttling^128^. This is in agreement with ΔNLS TDP-43 depletion effects demonstrated in vitro^129,130^ and in vivo^131^. Similarly, acetylation of K82 or/and K84 or single mutations of (K → A or ***R***) within the minor NLS, but not K95 (major NLS) resulted in impaired TDP-43 nuclear import and cytoplasmic mislocalization and aggregation *in vitro*^101,132^. Notably, acetylation of TDP-43 K82/84, and ubiquitination at K79 were found in insoluble fractions isolated from ALS patients suggesting involvement in disease pathogenicity^133^. Perhaps not coincidentally, these residues are structurally close to R83 and provide stability to the mNLS^99,101,126,130^. To note, enhanced importins binding to the R83-residue alleviated TDP-43 nuclear import deficits in several C9orf72-ALS/FTD models^134^, suggesting an important mechanistic role of citR83 TDP-43, and strengthening the rationale that the irreversible citrullination disturbs nucleocytoplasmic trafficking and TDP-43 homeostasis, promoting TDP-43 cytoplasmic mislocalization and pathogenicity.

As arginine is essential for TDP-43 phase-separation dynamics^117,119,120,135^, we demonstrate that citrullination of Low Complexity Domain (LCD)^136,137^ favors pathological liquid-solid phase separation state (LSPS) in recombinant systems. Consistent with these findings, we demonstrated that citrullination of citR268/272/275 residues in the CTD region promoted accumulation of soluble TDP-43 low-molecular weight proteoforms, identified as granular puncta accumulated in the cytoplasm. It is therefore possible that leveraging the immunoreactivity of the citR TDP-43 antibodies, we uncovered the first look into the morphology and biochemical properties of LSPS condensates presented across mice and human brain tissue.

Given the findings of the present study, it is intriguing to hypothesize that citrullination of RRMs motifs facilitates irreversible segment unfolding and exposure of residues to more common PTMs as recently reported following TDP-43 acetylation in cellular models^101,138^. Intriguingly, the disruption of TDP-43 RRM motif binding to nuclear GU-rich RNA was recently proposed to be a deciding mechanism for aberrant and passive TDP-43 nuclear export, contributing to increased cellular TDP-43 cytoplasmic concentrations^139^ and formation of solid-like toxic aggregates^82^. Although the effects of citR RRM motifs on nucleocytoplasmic transport were not investigated here, it is noteworthy to highlight that citR191 TDP-43 immunoreactivity was largely displayed in a nuclear/ perinuclear localization pattern in TAR4/4 mice and human tissue, while citR165 and R191 drove a moderate TDP-43 insolubility profile *in vivo*. Notably, the implications of RRMs citrullination on RNA metabolism and splicing of TDP-43 targets were not investigated in this study, but warrants further investigations.

Lastly, identification of novel putative citR TDP-43 conformers *in vitro* and *in vivo* models, and the confidence of the citR TDP-43 antibodies to recognize unique TDP-43 conformations, led us to hypothesize that citR TDP-43 inclusions are a hallmark of neuropathological changes associated with TDP-43 progression in LATE-NC, with or without comorbid ADNC. Immunohistochemical imaging analysis revealed epitope-specific profile of citR83 (NLS), citR191 (RRM2) and citR268/272 TDP-43 (LCD) in pure LATE-NC and ADNC+LATE-NC pathological progression and compared to the conventional pTDP-43 hallmark (Supplementary Table 3, Supplementary Fig. 5).

Collectively, our findings demonstrate for the first time the initial staging of citR TDP-43 intensities along LATE-NC and AD + LATE-NC pathologies and highlight the potential impact of citR on the clinically crucial stage 2 of LATE - NC^140,141^ (compiled in Supplementary Table 2). Importantly, our analysis provided the first insights on the contribution of citrullination on TDP-43 in human disease and predicted of citR TDP-43 conformers pathogenicity. We identified citR83 (NLS motif) and citR268/272 (CTD) as two consequential epitopes – able to drive formation of novel and heterogenous citR TDP-43 proteoforms in disease. Future and in-depth studies are crucial to unravel the mechanism(s) of epitope-specific citrullination on TDP-43 folding, RNA metabolism and the interplay with other PTMs and in synergy with proteinopathies (tau) as we recently reported ^81^.

## Acknowledgment

This research was supported by NIH R01AG084670-01/NIA (M-L.B.S), 1RF1AG072728/NIA (D.C.L), University of Kentucky Alzheimer’s Disease Research Center P30AG072946/NIA (L.V.E, M-L.B.S), NIH R01 NS105874/NINDS (A.C.M.F), Welch Foundation Q-2097-20220331 (J.C.F) and University of Kentucky, Sanders-Brown Start-Up funds (M-L.B.S).

## Author Contributions

Conceptualization, M-L.B.S, D.C.L; methodology, M-L.B.S., C.S., P.R.R., D.C., Z.Q., R.D., H.L., J.B.H., H.L., C.R., K.N., J.C.F., V.N.U.; investigation, C.S., P.R.R., R.D., Z.Q., H.L., J.B.H., H.L., C.R., K.N., E.L.A., C.N., P.S.T., V.N.U., P.T.N., J.C., D.C., D.C.L., M.-L.B.S.; resources D.C.L, A.C.M.F and P.T.N.; visualization, C.S., P.R.R., D.C., R.D., and V.N.U.; writing - original draft, M-L.B.S., C.S.,; writing - review & editing, M-L.B.S., C.S., P.T.N., and D.C.L; funding acquisition, M-L.B.S., and D.C.L.

## TABLES

**Table 1. Citrullinated TDP-43 residues and motif function in the C-terminal.** Mass spectrometry analysis identified five arginine epitopes citrullinated by PAD2 and PAD4 located within the C-terminal of TDP-43. Experiments were repeated twice.

**Table 2. Targeted citR antibody development strategy.** The peptide sequence and regional localization of citR epitopes targeted for generation of citR TDP-43 antibodies. Functional role of each region-containing citR TDP-43 epitope(s) is also indicated.

## SUPLEMENTARY TABLES

**Supplementary Table 1.** Identification of PAD-mediated TDP-43 citrullination. Mass spectrometry analysis identified eleven arginine epitopes citrullinated by PAD2 and PAD4. Peptide sequence of each citrullinated arginine and intensities are indicated for PAD2 and PAD4 reactions. Localization probability was determined as a value between 0 and 1 and posterior error probability values (PEP) for PADs - mediated activity are indicated.

| Loc. prob | PEP | Mass Error (ppm) | Arg (R) position | Sequence | Intensity |  |
| --- | --- | --- | --- | --- | --- | --- |
|  |  |  |  |  | PAD2 | PAD4 |
| 1 | 4.92e-39 | -0.00135 | R44 | LRYR(1)NPVSQC |  |  |
| 1 | 1.92e-37 | -0.18572 | R52 | MR(1)GVRLVEGI | 1.4e+07 | 3.4e+08 |
| 1 | 2.79e-21 | -0.30149 | R83 | NKR(1)KMDETDA | 4.7e+07 | 1.6e+08 |
| 1 | 2.20e-11 | -0.13541 | R165 | VMSQR(1)HMIDG | 2.6e+07 | 7.9e+07 |
| 1 | 3.11e-12 | 0.40841 | R171 | R(1)WCDCKLPNS | 3.3e+07 | 1.7e+07 |
| 1 | 7.70e-39 | 0.26697 | R189 | KQSQDEPLR(1)S | 9.3e+08 | 2.9e+08 |
| 1 | 6.71e-55 | 0.26697 | R191 | R(1)KVFVGRCTE | 6.2e+08 | 7.2e+08 |
| 1 | 5.93e-44 | -0.34384 | R268 | EPKHNSNR(1)QL | 3.1e+07 | 1.9e+08 |
| 1 | 5.93e-44 | 0.45386 | R272 | ER(0.99)SGRFGGNP | 3.9e+07 | 2.4e+08 |
| 1 | 1.31e-14 | 0.66957 | R275 | ERSGR(0.09)FGGNP | 2.8e+07 | 5.5e+06 |
| 1 | 7.76e-270 | -0.74837 | R293 | NSR(1)GGGAGLG | 9.2e+08 | 5.2e+08 |

**Supplementary Table 2.**
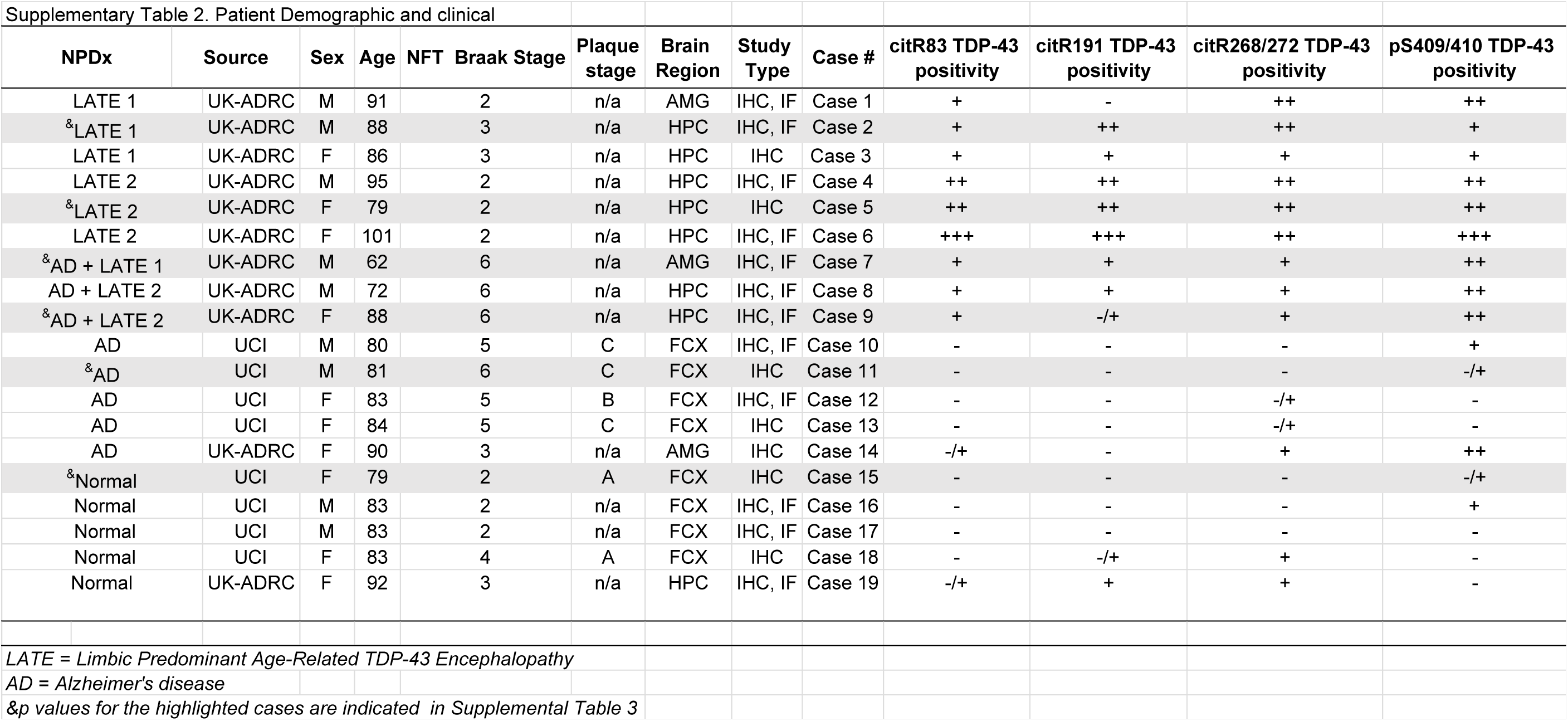
Demographics and clinical presentation. Demographics and pathologic features of the autopsy cohort cases received from the University of Kentucky Alzheimer’s Disease Research Center (UK-ADRC) and University of California, Irvine (UCI) brain bank and utilized in this study. Mean age at death for all LATE-NC stage 1 cases (n = 3) was 88.3 ± 2.05-year-old, LATE-NC stage 2 cases (n = 3) were 91.7 ± 9.29-year-old, AD + LATE-NC stage 1 + 2 (n = 3) was 74.0 ± 3.50 year old, mean age for AD cases (n = 5) was 83.6 ± 10.71-year-old and Healthy Control cases (n = 5) mean age was 84.0 ± 4.29 year old. Sex was indicated per each case. Neurofibrillary tangle (NFT) burden via Braak staging was determined during autopsy in all AD cases^67^. LATE staging was determined during autopsy and based on the consensus criteria^6^. *LATE – Limbic Predominant Age-Related TDP-43 Encephalopathy* *ADNC -* Alzheimer’s disease neuropathologic changes *- Not present, + Mild positivity, + + Moderate positivity, +++. High positivity*. *^&^ p values for the highlighted cases are indicated in Supplementary Table 3*

**Supplementary Table 3.**
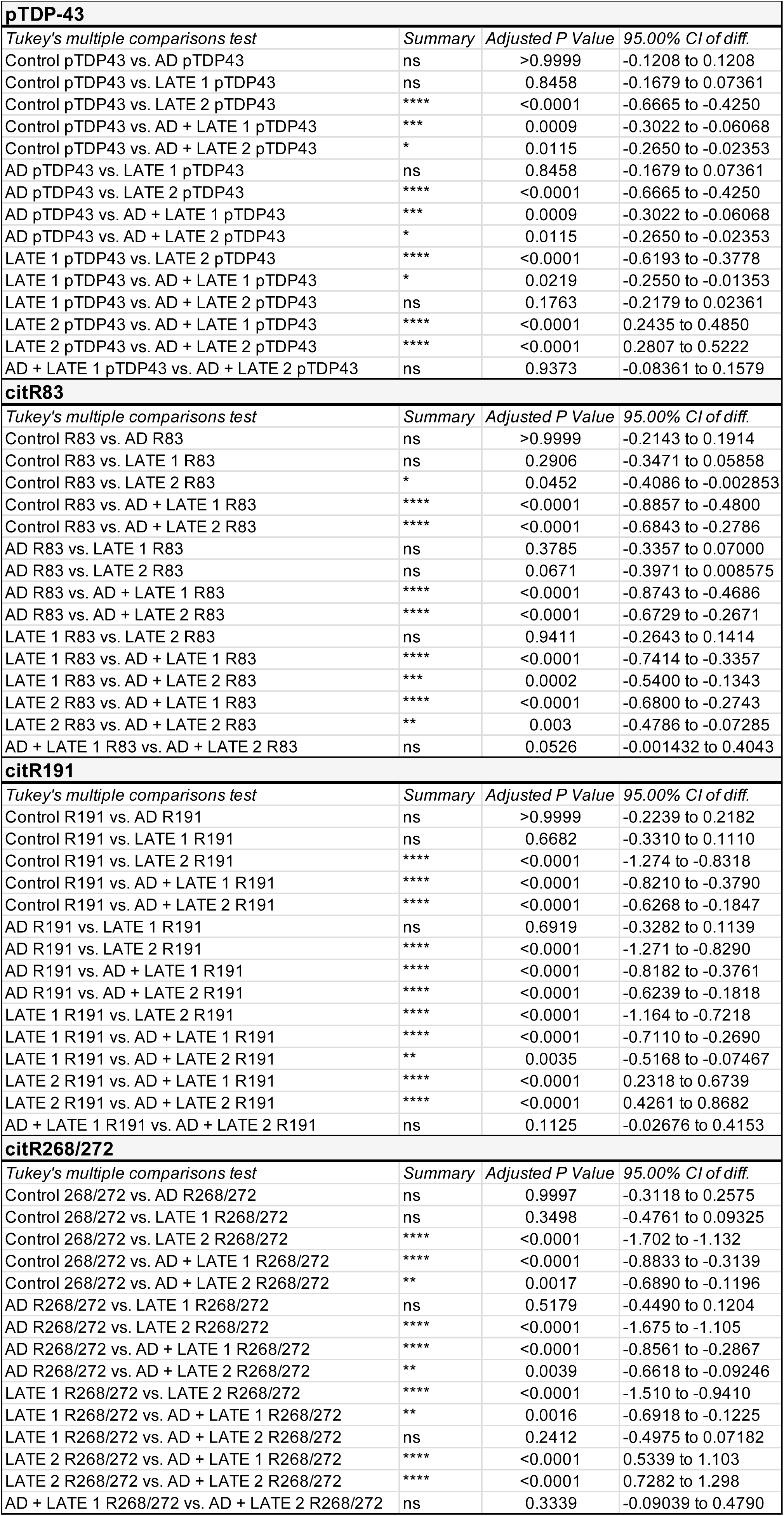
Statistical analysis of the neuropathological profile of citR TDP-43 in LATE-NC and ADNC + LATE-NC disease progression. Regional ROI (n = 7) intensity for citR TDP43 species and pTDP-43 in brain regions presented in each disease state (n = 1-2/ disease). Data represent the mean of the values ± SEM; One-way ANOVA, followed by Tukey’s post-hoc multiple comparisons tests, *p < 0.05, **p < 0.01, ***p < 0.001.

|  |  |  |  |
| --- | --- | --- | --- |
| **pTDP-43** |  |  |  |
| *Tukey's multiple comparisons test* | *Summary* | *Adjusted P Value* | *95.00% CI of diff.* |
| Control pTDP43 vs. AD pTDP43 | ns | >0.9999 | -0.1208 to 0.1208 |
| Control pTDP43 vs. LATE 1 pTDP43 | ns | 0.8458 | -0.1679 to 0.07361 |
| Control pTDP43 vs. LATE 2 pTDP43 | **** | <0.0001 | -0.6665 to -0.4250 |
| Control pTDP43 vs. AD + LATE 1 pTDP43 | *** | 0.0009 | -0.3022 to -0.06068 |
| Control pTDP43 vs. AD + LATE 2 pTDP43 | * | 0.0115 | -0.2650 to -0.02353 |
| AD pTDP43 vs. LATE 1 pTDP43 | ns | 0.8458 | -0.1679 to 0.07361 |
| AD pTDP43 vs. LATE 2 pTDP43 | **** | <0.0001 | -0.6665 to -0.4250 |
| AD pTDP43 vs. AD + LATE 1 pTDP43 | *** | 0.0009 | -0.3022 to -0.06068 |
| AD pTDP43 vs. AD + LATE 2 pTDP43 | * | 0.0115 | -0.2650 to -0.02353 |
| LATE 1 pTDP43 vs. LATE 2 pTDP43 | **** | <0.0001 | -0.6193 to -0.3778 |
| LATE 1 pTDP43 vs. AD + LATE 1 pTDP43 | * | 0.0219 | -0.2550 to -0.01353 |
| LATE 1 pTDP43 vs. AD + LATE 2 pTDP43 | ns | 0.1763 | -0.2179 to 0.02361 |
| LATE 2 pTDP43 vs. AD + LATE 1 pTDP43 | **** | <0.0001 | 0.2435 to 0.4850 |
| LATE 2 pTDP43 vs. AD + LATE 2 pTDP43 | **** | <0.0001 | 0.2807 to 0.5222 |
| AD + LATE 1 pTDP43 vs. AD + LATE 2 pTDP43 | ns | 0.9373 | -0.08361 to 0.1579 |
| **citR83** |  |  |  |
| *Tukey's multiple comparisons test* | *Summary* | *Adjusted P Value* | *95.00% CI of diff.* |
| Control R83 vs. AD R83 | ns | >0.9999 | -0.2143 to 0.1914 |
| Control R83 vs. LATE 1 R83 | ns | 0.2906 | -0.3471 to 0.05858 |
| Control R83 vs. LATE 2 R83 | * | 0.0452 | -0.4086 to -0.002853 |
| Control R83 vs. AD + LATE 1 R83 | **** | <0.0001 | -0.8857 to -0.4800 |
| Control R83 vs. AD + LATE 2 R83 | **** | <0.0001 | -0.6843 to -0.2786 |
| AD R83 vs. LATE 1 R83 | ns | 0.3785 | -0.3357 to 0.07000 |
| AD R83 vs. LATE 2 R83 | ns | 0.0671 | -0.3971 to 0.008575 |
| AD R83 vs. AD + LATE 1 R83 | **** | <0.0001 | -0.8743 to -0.4686 |
| AD R83 vs. AD + LATE 2 R83 | **** | <0.0001 | -0.6729 to -0.2671 |
| LATE 1 R83 vs. LATE 2 R83 | ns | 0.9411 | -0.2643 to 0.1414 |
| LATE 1 R83 vs. AD + LATE 1 R83 | **** | <0.0001 | -0.7414 to -0.3357 |
| LATE 1 R83 vs. AD + LATE 2 R83 | *** | 0.0002 | -0.5400 to -0.1343 |
| LATE 2 R83 vs. AD + LATE 1 R83 | **** | <0.0001 | -0.6800 to -0.2743 |
| LATE 2 R83 vs. AD + LATE 2 R83 | ** | 0.003 | -0.4786 to -0.07285 |
| AD + LATE 1 R83 vs. AD + LATE 2 R83 | ns | 0.0526 | -0.001432 to 0.4043 |
| **citR191** |  |  |  |
| *Tukey's multiple comparisons test* | *Summary* | *Adjusted P Value* | *95.00% CI of diff.* |
| Control R191 vs. AD R191 | ns | >0.9999 | -0.2239 to 0.2182 |
| Control R191 vs. LATE 1 R191 | ns | 0.6682 | -0.3310 to 0.1110 |
| Control R191 vs. LATE 2 R191 | **** | <0.0001 | -1.274 to -0.8318 |
| Control R191 vs. AD + LATE 1 R191 | **** | <0.0001 | -0.8210 to -0.3790 |
| Control R191 vs. AD + LATE 2 R191 | **** | <0.0001 | -0.6268 to -0.1847 |
| AD R191 vs. LATE 1 R191 | ns | 0.6919 | -0.3282 to 0.1139 |
| AD R191 vs. LATE 2 R191 | **** | <0.0001 | -1.271 to -0.8290 |
| AD R191 vs. AD + LATE 1 R191 | **** | <0.0001 | -0.8182 to -0.3761 |
| AD R191 vs. AD + LATE 2 R191 | **** | <0.0001 | -0.6239 to -0.1818 |
| LATE 1 R191 vs. LATE 2 R191 | **** | <0.0001 | -1.164 to -0.7218 |
| LATE 1 R191 vs. AD + LATE 1 R191 | **** | <0.0001 | -0.7110 to -0.2690 |
| LATE 1 R191 vs. AD + LATE 2 R191 | ** | 0.0035 | -0.5168 to -0.07467 |
| LATE 2 R191 vs. AD + LATE 1 R191 | **** | <0.0001 | 0.2318 to 0.6739 |
| LATE 2 R191 vs. AD + LATE 2 R191 | **** | <0.0001 | 0.4261 to 0.8682 |
| AD + LATE 1 R191 vs. AD + LATE 2 R191 | ns | 0.1125 | -0.02676 to 0.4153 |
| **citR268/272** |  |  |  |
| *Tukey's multiple comparisons test* | *Summary* | *Adjusted P Value* | *95.00% CI of diff.* |
| Control 268/272 vs. AD R268/272 | ns | 0.9997 | -0.3118 to 0.2575 |
| Control 268/272 vs. LATE 1 R268/272 | ns | 0.3498 | -0.4761 to 0.09325 |
| Control 268/272 vs. LATE 2 R268/272 | **** | <0.0001 | -1.702 to -1.132 |
| Control 268/272 vs. AD + LATE 1 R268/272 | **** | <0.0001 | -0.8833 to -0.3139 |
| Control 268/272 vs. AD + LATE 2 R268/272 | ** | 0.0017 | -0.6890 to -0.1196 |
| AD R268/272 vs. LATE 1 R268/272 | ns | 0.5179 | -0.4490 to 0.1204 |
| AD R268/272 vs. LATE 2 R268/272 | **** | <0.0001 | -1.675 to -1.105 |
| AD R268/272 vs. AD + LATE 1 R268/272 | **** | <0.0001 | -0.8561 to -0.2867 |
| AD R268/272 vs. AD + LATE 2 R268/272 | ** | 0.0039 | -0.6618 to -0.09246 |
| LATE 1 R268/272 vs. LATE 2 R268/272 | **** | <0.0001 | -1.510 to -0.9410 |
| LATE 1 R268/272 vs. AD + LATE 1 R268/272 | ** | 0.0016 | -0.6918 to -0.1225 |
| LATE 1 R268/272 vs. AD + LATE 2 R268/272 | ns | 0.2412 | -0.4975 to 0.07182 |
| LATE 2 R268/272 vs. AD + LATE 1 R268/272 | **** | <0.0001 | 0.5339 to 1.103 |
| LATE 2 R268/272 vs. AD + LATE 2 R268/272 | **** | <0.0001 | 0.7282 to 1.298 |
| AD + LATE 1 R268/272 vs. AD + LATE 2 R268/272 | ns | 0.3339 | -0.09039 to 0.4790 |

## SUPLEMENTARY VIDEOS

**Supplementary Video 1. TDP-43 aggregate morphology.** Wild-type TDP-43 [5µM] incubated for 72 hours under 25⁰C, 800 CFM orbital rotation conditions. TEM tomography of the aligned cross-sectional TDP-43 aggregate images were processed with ImageJ for single model viewing using the aligned stack projection feature. The resulting stack was converted to .AVI format and exported in video form.

**Supplementary Video 2. Citrullinated TDP-43 condensate morphology.** and citR TDP-43 [5µM] were incubated for 72 hours under 25⁰C, 800 CFM orbital rotation conditions. TEM tomography of the aligned cross-sectional citR TDP-43 images were processed with ImageJ for single model viewing using the aligned stack projection feature. The resulting stack was converted to .AVI format and exported in video form.

**Supplementary Video 3.** Video production was generated using Fusion360 software (Autodesk) from the raw tomography binary stacks. citR TDP-43 size and condensate composition is shown in comparison to TDP-43 aggregates. Scale is conserved and kept consistent from raw tomography projections above (Supp. Video 1 and 2).

## SUPPLEMENTARY FIGURE LEGEND

**Supplementary Figure 1.**
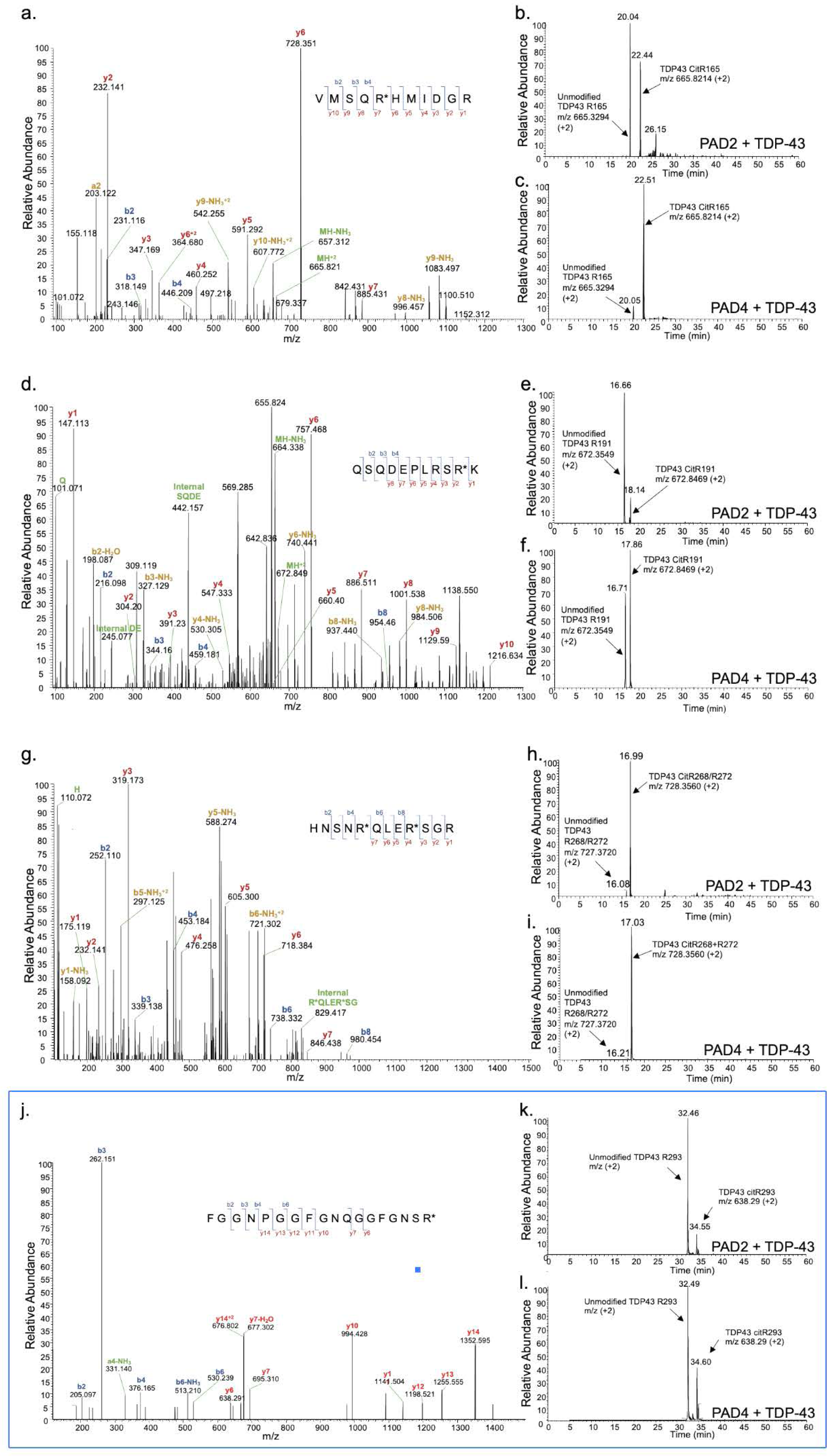
Identification of citrullinated TDP-43 epitopes. Spectra showing b- and y-ion coverage of citrullinated peptides, as well as XICs retention times for unmodified and citrullinated peptides treated with PAD2 and PAD4, corresponding to a - c, citR165 (intact peptide monoisotopic m/z (+2) 665.8214), d - f, citR191(intact peptide monoisotopic m/z (+2) 672.8469), g - i, citR268/272 (intact peptide monoisotopic m/z (+2) 728.3560), j - l, citR293 (intact peptide monoisotopic m/z (+2) 638.29).

**Supplementary Figure 2.**
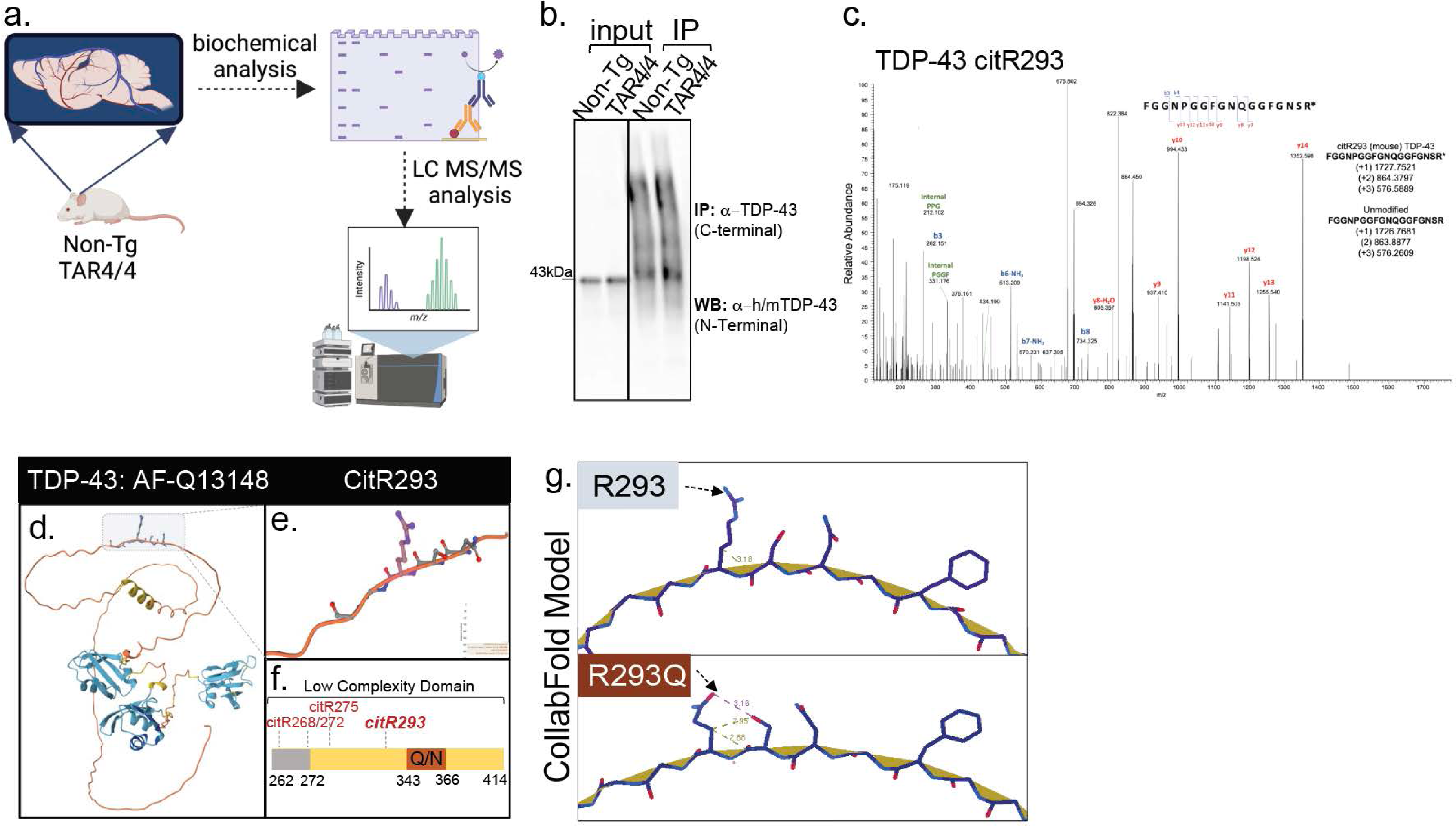
Identification of citR293 TDP-43 in vivo via IP-MS. a, Schematic rendering of assays identifying citR293 TDP-43 in the TAR4/4 mouse brain tissue. b, Images of the cortical brain homogenate (input) and immunoprecipitated fraction from TAR4/4 and Non-Tg mice using human/mouse TDP-43 (C-terminal) antibody. Western blot membrane was probed with a total TDP-43 antibody. c, MS/MS spectra of the b- and y-ion coverage of citrullinated R293 peptides identified in the IP cortical fraction of the TAR4/4 mice (n = 2). d, Alpha-fold renderings of TDP-43 (PDB ID:7KWZ, AF -Q13148) highlighting e, low complexity domain and sequence linearity rendering citR293 epitope at 5Å, f, Schematic of the LCD region of TDP-43 showing relative locations of three citR sites: citR268/272, citR275, and citR293. g, Schematic rendering of CollabFold folding prediction with R->Q theoretical mutagenesis.

**Supplementary Figure 3.**
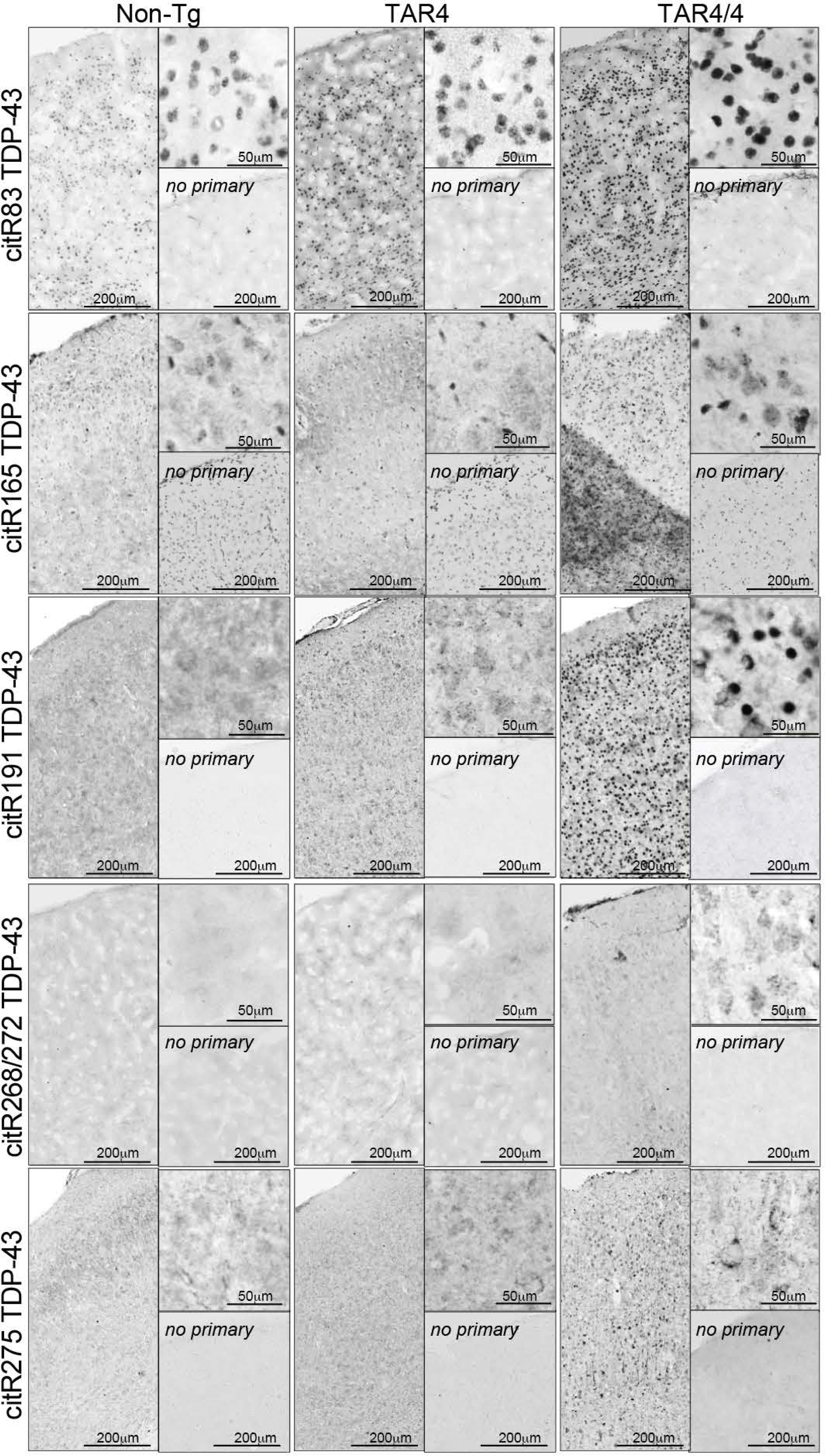
Citrullination of TDP-43 is induced in TDP-43 mouse models. a, Immunohistochemical images of cortical tissue labeled with citR83, 165, 191, 268/272, and 275, TDP-43 antibodies. High magnification images of labeled neurons and no primary controls are also shown. Scale bar is 50µm and 200 µm.

**Supplementary Figure 4.**
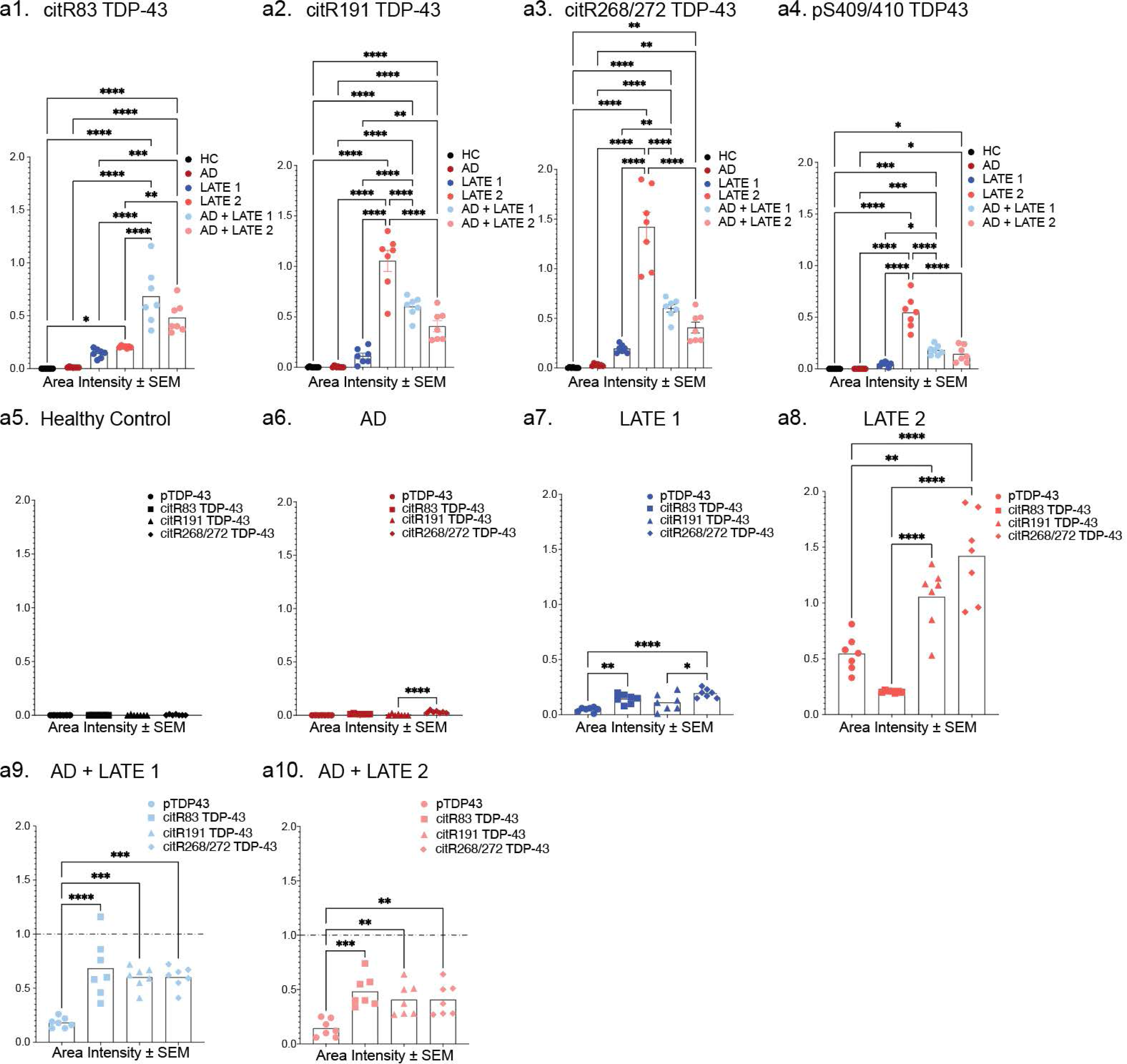
Analysis of citrullinated TDP-43 levels in ADNC / LATE-NC disease states. Regional ROI (n = 7) intensity ratio to HC of each citR TDP43 species and pTDP-43 in brain regions presented in each disease state (n = 1 - 2). a1 - a4, inter-disease analysis for each citR epitope and a5 - a10, intra-citR TDP-43 intensity relationship is analyzed by GraphPad. Data represent the mean of the values ± SEM; One-way ANOVA, followed by Tukey’s or Šidák *post hoc* multiple comparisons tests, *p < 0.05, **p < 0.01, ***p < 0.001.

**Supplementary Figure 5.**
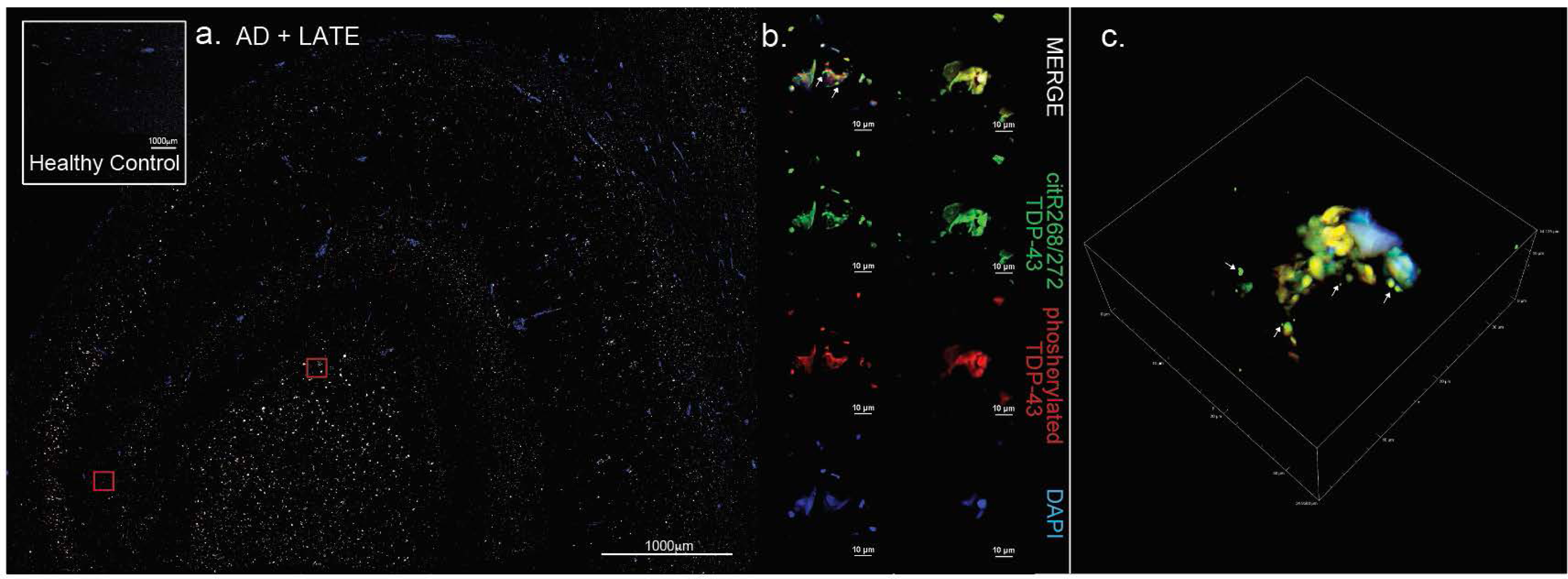
Co-localization of citR83 TDP-43 with pTDP-43 in ADNC + LATE-NC stage 2. Immunofluorescence (IF) images of hippocampal tissue from an AD + LATE stage 2 case vs. HC (inset) labeled against citR83 (AF488) and pTDP-43 (AF594). b, magnified images for each modification and merged images demonstrated co-localization between citR and pTDP-43 species, while singly labeled citR83 (+) condensates were also observed (white arrows). c, 3D max intensity projection of merged images. All images taken by confocal z-stack, 20x magnification with 2x NyQuist optical zoom, resonant scan with sequential channel imaging, large image stitch of entire tissue.

**Supplementary Figure 6.**
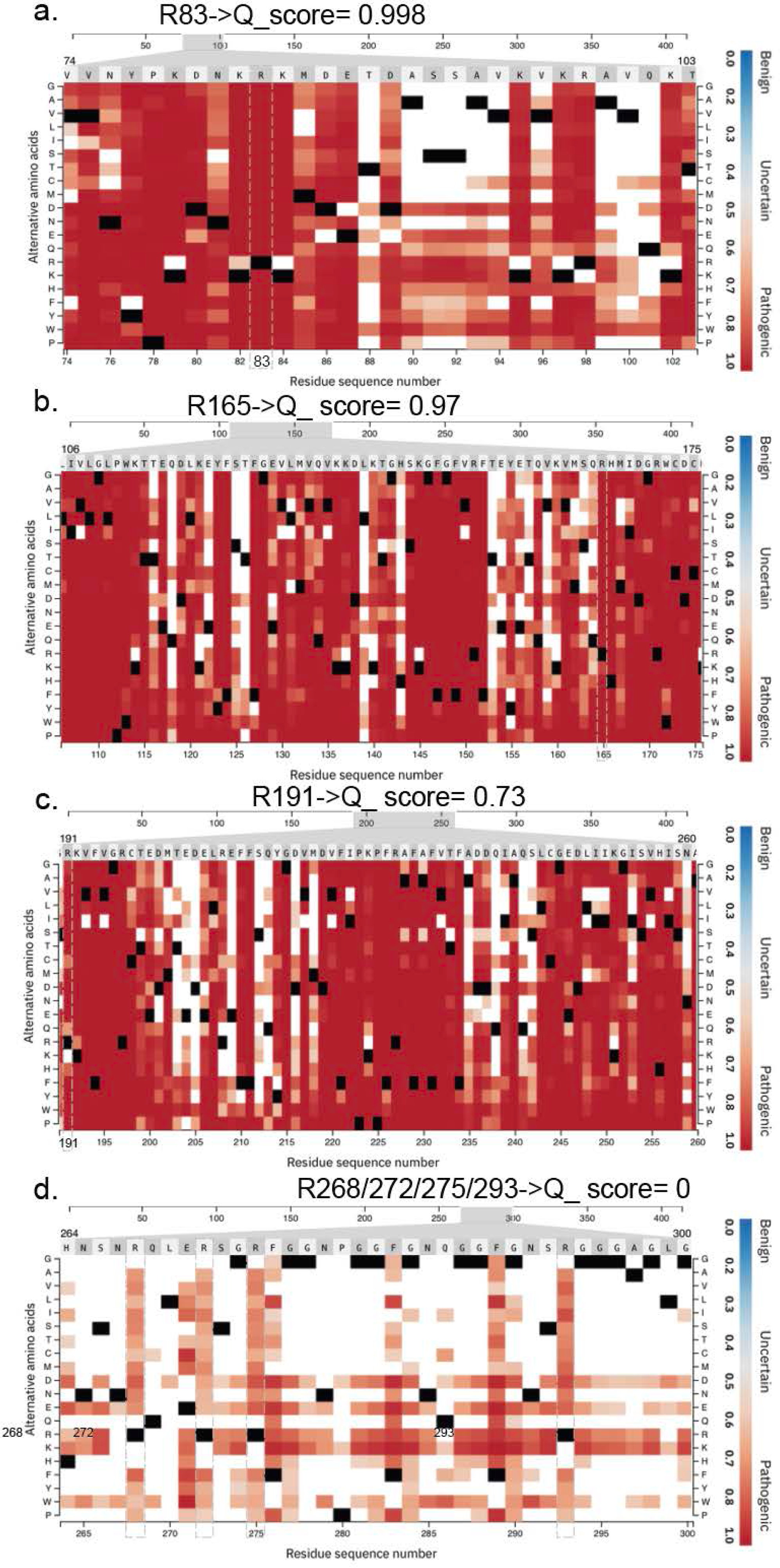
Pathogenicity of AlphaMissense R -> Q TDP-43 “citrullination mimics”. Pathogenicity of each confirmed citrullinated arginine within TDP-43 functional domains analyzed in AlphaFold2. Citrullination mimic of a, R83 → Q pathogenetic score (0.998) within the NLS domain (position range= 0.986 - 1). b, R165 → Q pathogenetic score (0.978) within the RRM1 (position range= 0.917 - 1). c, R191 → Q pathogenetic score (0.737) within the RRM2 domain (position range= 0.737 - 0.998). d, Single citrullination mimics of R268/272/275/293 → Q did not yield a pathogenicity score within the CTD region (position range= 0.626 - 0.860 *vs.* 0.561 - 0.838 *vs.* 0.643 - 0.885 *vs.* 0.643 - 0.894, respectively).

## Abbreviations

PAD: peptidyl arginine deiminase
PTM: post-translational modification
TDP-43: TAR DNA Binding Protein 43 kDa
CitR: Citrullinated
LATE: Limbic predominant Age-related TDP-43 Encephalopathy
AD: Alzheimer’s disease
ALS: Amyotrophic lateral sclerosis
FTD: Frontotemporal dementia
RFU: relative fluorescent units
LLPS: liquid-liquid phase separation
LSPS: liquid-solid phase separation

